# Disruption of Cell-Type-Specific Molecular Programs of Medium Spiny Neurons in Autism

**DOI:** 10.1101/2025.11.05.686845

**Authors:** Guohua Yuan, Varun Suresh, Emilie Wigdor, Yuhan Hao, Rachel Leonard, Marilyn Steyert, Michael Griffiths, Clements Evans, Narjes Rohani, Jakob Weiss, Frederik H. Lassen, Nicole Schafer, Shan Dong, Duncan S. Palmer, Stephan J. Sanders, Tomasz J. Nowakowski

**Affiliations:** Department of Neurological Surgery, University of California San Francisco, San Francisco, CA, USA; Institute of Developmental and Regenerative Medicine, Department of Paediatrics, University of Oxford, Oxford, UK; Sir William Dunn School of Pathology, University of Oxford, Oxford, UK; Center for Human Genetics, University of Oxford, Oxford, UK; The Pioneer Centre for SMARTbiomed, Big Data Institute, Li Ka Shing Centre for Health Information and Discovery, University of Oxford, Oxford, UK; Department of Psychiatry and Behavioral Sciences, University of California San Francisco, San Francisco, CA, USA; Weill Institute for Neurosciences, University of California San Francisco, San Francisco, CA, USA; Program in Medical and Population Genetics, Broad Institute of MIT and Harvard, Cambridge, MA, USA; Department of Statistics, University of Oxford, Oxford, UK; New York Genome Center, New York, NY, USA; Department of Anatomy, University of California San Francisco, San Francisco, CA, USA; Eli and Edythe Broad Center for Regeneration Medicine and Stem Cell Research, University of California San Francisco, San Francisco, CA, USA

## Abstract

Autism spectrum disorders (ASD) are highly heritable neurodevelopmental conditions with major contributions from rare genetic variants. Most studies have focused on cortical mechanisms; even growing evidence implicates subcortical circuits in ASD etiology. To systematically map developmental and molecular alterations beyond the cortex, we profiled lineage relationships across five brain regions in an ASD mouse model. Most prominent changes emerged in the striatum, a hub for learning and motor control. Furthermore, we performed single-nucleus multiomic profiling of human putamen from ASD and neurotypical donors revealed cell-type-specific transcriptomic and regulatory alterations. Differential expression converged on synaptic and energy metabolic dysfunctions in D1 striosome medium spiny neurons (MSNs), coupled with astrocytic remodeling of synaptic support. Gene regulatory network analysis identified EGR3 and EGR1 as key transcriptional regulators of ASD-associated programs of D1 MSNs. Together, these results establish the striatum as a central node of ASD convergence and provide a multiomic resource for dissecting its subcortical mechanisms.

## Introduction

Autism spectrum disorder (ASD) is a complex neurodevelopmental condition characterized by persistent deficits in reciprocal social communication, alongside restricted and repetitive patterns of behavior, interests, or activities^1,2^. ASD is highly heritable, with twin-based heritability estimates exceeding 70%^3,4^. *De novo* gene disrupting variants contributing to ASD in ∼13% of individuals have been identified^5,6^. Among these, the recurrent ∼0.7 Mb deletion at 16p11.2 is one of the most common and largest single genetic influences associated with ASD^7–9^.

While genetic factors such as the recurrent 16p11.2 deletion provide key insights into ASD etiology, investigations at the neural systems level have identified various structural and functional abnormalities across multiple brain regions in ASD. Nevertheless, most transcriptomic and epigenomic investigations have primarily focused on cortical tissues^10–12^. This cortical focus has left substantial gaps in our understanding of subcortical contributions to ASD pathophysiology, despite mounting evidence for subcortical involvement in the condition^13–19^.

To address this critical knowledge gap, we first performed an unbiased lineage-tracing screen across five brain regions (cortex, hippocampus, striatum, thalamus, olfactory bulb) in a 16p11.2 microdeletion mouse model, which pinpointed the striatum as the principal locus of change, with strong shifts in cell-fate composition and differential gene expression within medium spiny neuron (MSN) subtypes. Initial studies suggest ASD-associated genes are enriched in striatal medium spiny neurons^13,20^, however, comprehensive molecular characterization of cell identity and fate changes in subcortical regions, such as the putamen, remains limited in ASD.

The striatum plays a pivotal role in motor control, reward processing, and cognitive flexibility; functions often disrupted in ASD^17,21,22^. As the major component of the dorsal striatum, the putamen receives extensive projections from cortical and thalamic regions and serves as a key node linking social deficits and repetitive behaviors in autism pathophysiology ^15^. Neuroimaging studies have revealed structural and functional alterations in the putamen in ASD, including significantly altered volumes and atypical connectivity patterns with cortical regions^15,23,24^. Furthermore, repetitive behaviors in autism are specifically linked to putaminal dysfunction and imbalanced corticostriatal connectivity^25^.

To determine if dysregulation of striatal cell types could also take place in humans, we performed single-nucleus multiomic profiling of putamen postmortem tissue from 84 donors (37 neurotypical and 47 ASD), through ages 0-49; thus making our cohort of donors one of the largest known single cell mapping repository of the human putamen. Mapping cell type–specific transcriptomic alterations revealed that differentially expressed genes (DEGs) were predominantly converged in D1 MSNs. These DEGs converged on synaptic pathways, indicating reduced synaptic efficacy and impaired energy metabolism that may render D1 striosome neurons more vulnerable to stress, resembling early stages of neuronal degeneration. In ASD, altered gene programs also exhibited signs of astrocytic remodeling of synaptic support,and oligodendrocytic perturbations possibly affecting myelination. Integrative regulatory analyses combining chromatin accessibility and gene expression highlighted *EGR3* and *EGR1* as candidate transcriptional drivers organizing cell type–specific gene-regulatory networks. Together, these data show that cellular and molecular changes in a 16p11.2 microdeletion model and in humans with ASD with (N=14) and without (N=33) a genetic diagnosis, converge on the striatum, providing a reference multi-omic resource for the human putamen in ASD and a framework for dissecting subcortical mechanisms.

## Results

### Systematic evaluation of lineage and transcriptomic differences across five brain regions in 16p11.2df mice

The proximal 16p11.2 deletion (BP4-BP5, chr16:29,639,519-30,189,452 hg38) represents one of the most prevalent genetic copy number variants (CNVs) associated with ASD^26^. How this mutation interferes with neurodevelopmental processes of cell fate specification has not been systematically investigated. Recent studies suggest that subcortical structures may show stronger transcriptional responses to large CNV microdeletions^27,28^, but most transcriptomic studies have focused on cortical tissues^29,30^.

To survey clonal cell lineage relationships in a genetic mouse model of ASD carrying defined genetic alterations, we leveraged “scRNA-seq-compatible tracer for identifying clonal relationships” (STICR)^31^ to conduct combinatorial barcoding of individual progenitor cells of mouse embryos at E12.5 for both 16p11.2df (#013128 : B6129S-Del(7Slx1b-Sept1)4Aam/J) and wild-type mice. We harvested the brains at postnatal day 4 (P4), and dissected the tissue to enrich cells in the cortex (rostral and caudal separated), striatum, thalamus, hippocampus, and the olfactory bulb (OB) (Figure 1A). We recovered reporter expressing cells using fluorescence activated cell sorting and processed the cells for scRNAseq (see Methods) to recover transcriptional profiles along with clonal cell lineage relationships. With error-correction, STICR enables clonal lineage tracing of up to 250,000 individual cells per experiment, with a barcode collision probability of less than 0.5%.

**Figure 1.**
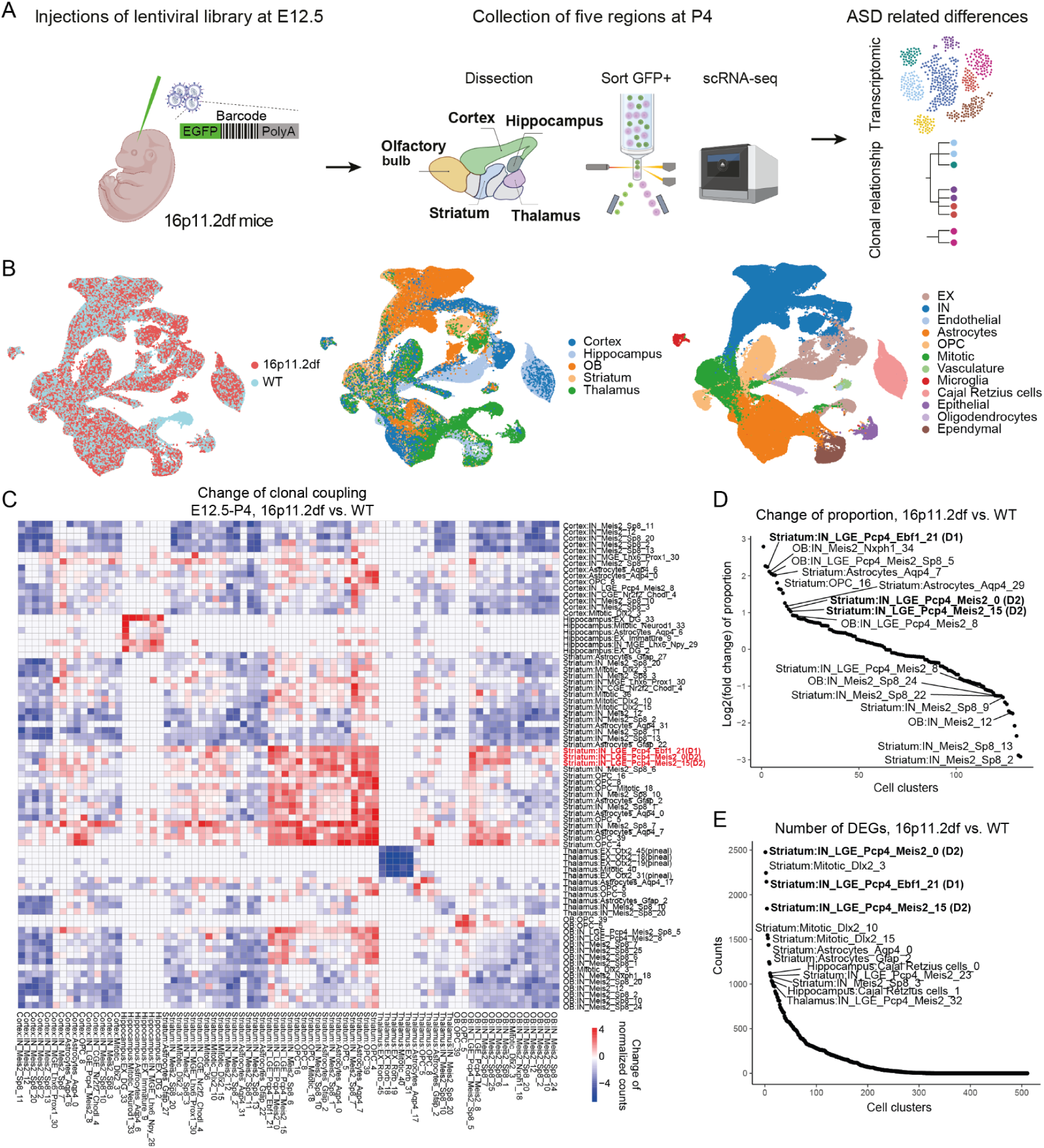
Clonal and transcriptional differences in multiple brain regions of 16p11.2df mice. (A) Schematic diagram illustrating the workflow for data collection. (B) UMAP visualization of cells collected from 16p11.2df and wild-type mice, labeled by viral injections at E12.5 and collected at P4. Cells are colored by genotype (left), regions (middle), and annotated cell types (right). (C) Heatmap showing the difference in clonal coupling of cell clusters across all regions between 16p11.2df and wild-type mice injected at E12.5. Colors represent changes in clonal coupling, calculated as the difference between normalized shared clonal counts. (D) Fold change of cell type proportions in each sub-cell cluster between 16p11.2df and wild-type mice injected at E12.5. Cells in multi-cellular clones are included in the analysis. Clusters with more than 0.1% cells in both phenotypes are included in the analysis. (E) Number of DEGs in each sub-cell cluster between 16p11.2df and wild-type mice injected at E12.5. Clusters with the highest numbers of DEGs are labeled.

We captured 635,000 cells and, after quality filtering (removing low-quality cells, potential doublets, and high-doublet ratio clusters), retained 269,499 high-quality cells with clonal barcodes detected, including 128,836 cells from WT and 140,613 cells from 16p11.2df samples. These cells were obtained from 14 brains across 57 10X reactions, with more than 40,000 cells per region (Table S1). We identified 157,600 unique barcodes, of which 45,300 contained multiple cells, which we henceforth refer to as multicellular clones (Table S2). The proportion of shared clones between different brains remained below 0.5%, validating the theoretical estimates based on the diversity of the viral library (Figure S1A).

We retained cells with a clonal barcode and transcriptome for further analysis (Figure 1B), starting with dimensionality reduction and unbiased clustering using the shared nearest neighbor (SNN) graph constructed from high-dimensional space. We annotated cell identities based on cluster-wise enriched expression of canonical marker genes (Figure 1B, Figure S1B). We identified glutamatergic neurons (*Slc17a7*, *Slc17a6*, *Neurod2*), GABAergic neurons (*Dlx2*, *Gad1*, *Gad2*), oligodendrocyte progenitor cells (OPC) (*Olig1*, *Olig2*, *Pdgfra*), oligodendrocytes (*Mbp*, *Mog*), astrocytes (*Gfap*, *Aqp4*, *Apoe*), Cajal–Retzius (CR) (*Reln*) cells, Ependymal cells (*Foxj1*), choroid plexus epithelial cells (*Ttr*), endothelial cells (*Esam*), microglia (*Ctss*), and vascular cells (*Vtn*). We further identified subtypes of glutamatergic neurons based on distinct molecular markers of each region, such as upper cortical layer 2 and 3 (L2/3, *Cux1*, *Cux2*), layer 4 (L4, *Rorb*), and deep cortical layer 5 and 6 (L5/6, *Bcl11b*; *Scg2*), CA1 (*Mapped1*), and dentate gyrus (DG) (*Prox1*) neurons of the hippocampus. GABAergic neurons can be subdivided into several cardinal classes according to the neurotransmitter character (e.g., *Vip*, *Sst*), or based on the potential progenitor domain of origin. Cell type annotation was performed on wild type data and then the 16p11.2df single-cell data was projected into the wild-type reference and annotated by label transferring. Consistent with the expected cell type proportions, striatum, and OB samples were enriched for GABAergic neurons, while the hippocampus was largely composed of glutamatergic neurons^32,33^ (Figure S1C).

To study the regional specific lineage progression patterns and the regional dispersed clones, we calculated the clone coupling between different cell types in each region by the normalized number of shared clones between each other. To determine altered cell lineage relationships, we compared clonal coupling between cluster pairs by subtracting normalized clonal coupling counts between 16p11.2df and wild type (Figure 1C, Figure S2A). We identified differences in clonal relations involving 511 cluster pairs, including 13 broad cell types found within and across brain regions. Strikingly, the largest relative number of cell types whose clonal cell lineage relations were significantly altered were found within the striatum (Figure 1C).

Most notably, clonal relationships of D1 (cluster 21) and D2 (clusters 0 and 15) medium spiny neurons (MSNs) showed pronounced alterations in the striatum (Figure 1C). D1 and D2 MSNs are often clonally related and generated from shared LGE progenitors during mid-gestation, with D1 neurons produced earlier (E12.5–E14.5) and D2 neurons emerging slightly later (E13.5–E16.5) during normal striatal development^34–36^. Both D1 and D2 neurons exhibited an increased number of clones in the 16p11.2df dataset, along with astrocytes and OPCs from the same lineage (Figure 1C–1D). Notably, the changes were more pronounced in D1 neurons than in D2 neurons. Cell type–specific differential gene expression analysis was performed at the cluster level to identify transcriptional differences between 16p11.2df and wild-type mice. The differentially expressed genes (DEGs) found in D1 and D2 neurons (N= 3590 unique genes) ranked among the highest across all clusters (Figure 1E). Together, these findings highlight distinct spatial-temporal disruptions in clonal relationships, suggesting that the 16p11.2df genetic variant influences specific neuronal lineages and transcriptional programs, most notably in subcortical regions, such as striatum, which are often understudied.

### Single-nucleus multi-omic atlas of human ASD and neurotypical putamen

To investigate if any of the changes identified in a genetic mouse model extend to humans, we generated single-nucleus RNA/ATAC-seq data from the putamen of 84 donors obtained from the Autism BrainNet bank (Table S3). The cohort included 37 controls without known neuropsychiatric disorders, 47 autistic individuals, aged 1 to 49 years, of whom 33 without a known genetic diagnosis, and 14 individuals with a neurodevelopmental disorder (NDD) and an ASD-associated genetic diagnosis (Figure 2A) of genetic variants: 15q11.2-13.1 duplication, 15q11.2-13.1 deletion, 22q13.3 deletion (*SHANK3*), *KMT2A*, *GRIN2B*, *STXBP1*, *ANK2*, and *KIF1A*. To minimize batch effects and avoid confounding with case status, we pooled tissues from ∼10-12 individuals (∼50% case or control) as a group to perform multiple (∼4 per tissue group) single-nucleus multiome (snMultiome) 10x Genomics assays (Figure 2A and Figure S4A). We then demultiplexed donors by matching genotypes of common variants called from snRNA and genotype array data using demuxlet (see Methods). We initially recovered 418,525 nuclei and obtained 346,556 paired single-nucleus RNA-seq and ATAC-seq data after quality control (Figure S3). On average we detected 2,019 genes, 5,352 transcripts, and 14,811 ATAC fragments per nucleus.

**Figure 2.**
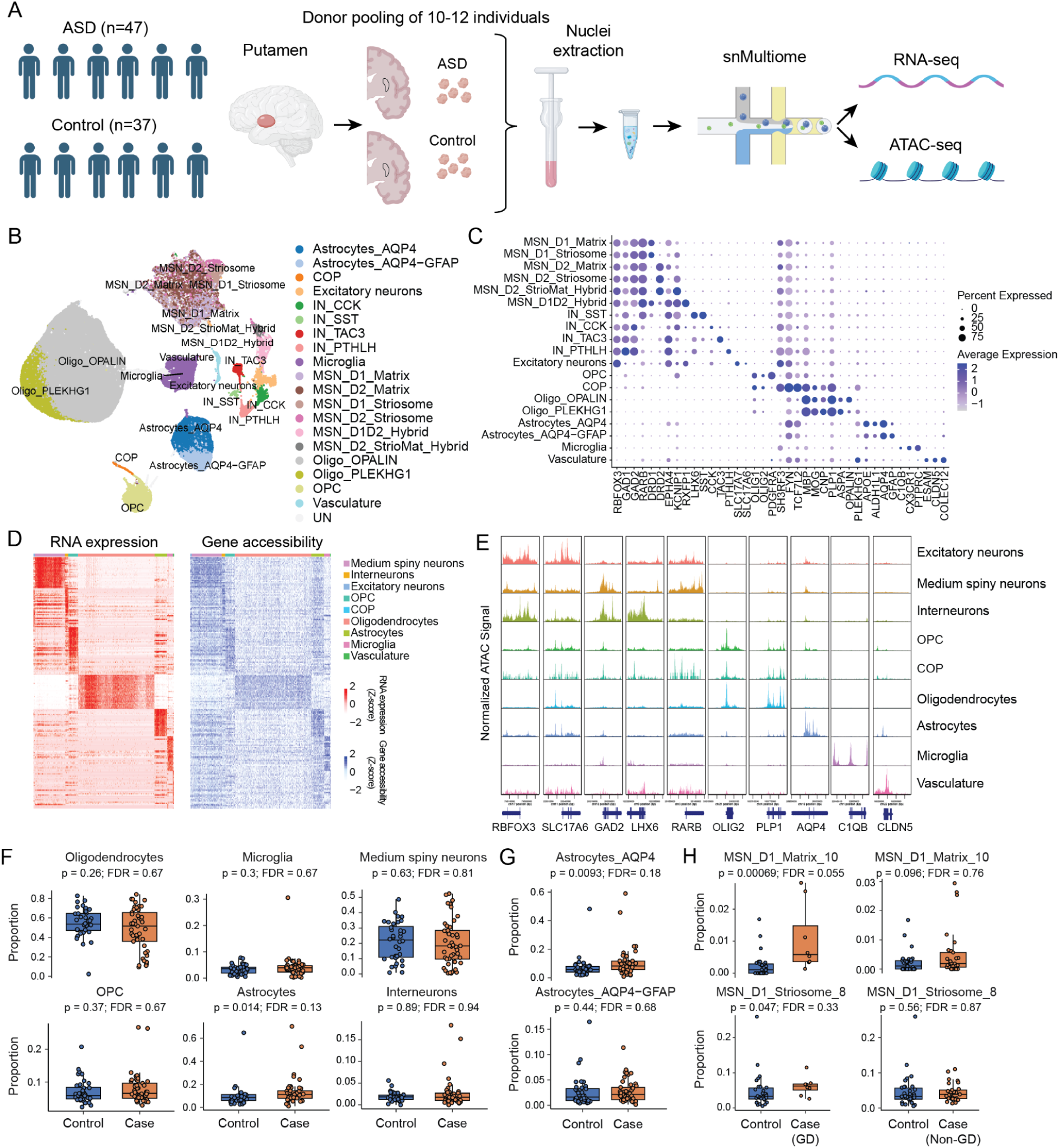
Single-nucleus multiomic profiling of human putamen in ASD and controls. (A) Schematic overview of the study design. Tissues from donors with ASD and controls were pooled as a tissue group prior to nuclei extraction to minimize batch effects between samples. Donor identities were subsequently demultiplexed using single-nucleotide polymorphism (SNP) array information derived from the sequencing reads of the single-cell multiome data. (B) UMAP visualization of high-quality nuclei colored by annotated cell type. MSN, medium spiny neuron; OPC, oligodendrocyte progenitor cell; COP, committed oligodendrocyte precursor. (C) Dot plot showing cell-type marker expression across annotated cell types. Dot size represents the percentage of nuclei expressing each gene, and color intensity denotes scaled average expression. (D) Heatmaps of z-score normalized RNA expression (left) and gene chromatin accessibility (right) showing their concordant changes across cell types. Each row represents a gene. Each column represents a cell. (E) Representative genome tracks displaying ATAC-seq signal at canonical marker gene loci. (F) Quantification of major cell-type proportions between ASD and control donors. The proportion was normalized to all cells. P values were obtained using the Wilcoxon test and FDR was calculated by using the BH method. (G) Subtype-level analysis reveals that the increase in astrocyte proportion is driven primarily by *AQP4*⁺ homeostatic astrocytes rather than *GFAP*⁺ reactive astrocytes. Proportions were normalized to the total cell population. P values were obtained using the Wilcoxon test and FDR was calculated by using the BH method. (H) MSN subtype composition analysis showing the increased cell type proportions in D1 MSN populations. The proportion was normalized to all MSNs. P values were obtained using the Wilcoxon test and FDR was calculated by using the BH method. GD: Cases with genetic diagnosis. Non-GD: Cases without genetic diagnosis.

We performed unsupervised clustering of single-cell transcriptomes and annotated cell types based on the expression of canonical marker genes (Figures 2B–2C and Figures S4 B-C). The major identified populations included medium spiny neurons (MSNs) (*GAD2*, *RARB*), interneurons (*GAD2*, *LHX6*), excitatory neurons (*SLC17A7*, *SLC17A6*), oligodendrocytes (*MBP*, *MOG*), oligodendrocyte progenitor cells (OPC) (*OLIG1*, *OLIG2*), committed oligodendrocyte precursor cells (COP) (*SH3FR3*, *FYN*), astrocytes (*GFAP*, *AQP4*), microglia (*CX3CR1*, *P2RY12*), and vascular cells (*ESAM*, *CLDN5, COLEC12*). Consistent with the known cellular composition of human striatum^37^, the dataset was enriched for medium spiny neurons and oligodendrocytes (Figure S4D). Within interneurons, we identified transcriptionally distinct subtypes, including *SST⁺*, *CCK⁺*, *TAC3⁺*, and *PTHLH⁺* populations. Each exhibits diverse functions and modulates the excitability and inhibition of MSNs in response to various neuromodulatory stimuli^38,39^. Oligodendrocytes were subdivided into *PLEKHG1⁺* and *OPALIN⁺* subtypes, corresponding to pre-myelinating and mature myelinating states, respectively. Among astrocytes, a subtype with enriched GFAP expression was identified, which is related to reactive astrocytes under inflammation^40^. To resolve MSN diversity, we mapped our dataset to the Cell Type Taxonomies of the human basal ganglia using MapMyCells (RRID:SCR_024672). This revealed distinct D1 (*DRD1*), D2 (*DRD2*), and D1/D2 hybrid (*RXFP1*) subtypes (Figure S4 E-F), including matrix (*EPHA4*), striosome (*KCNIP1*), and striosome/matrix hybrid MSNs (*CNTN5*), which differ in striatal compartmentalization, dopamine receptor expression, and projection targets^41^.

To identify marker genes enriched in each cell population, we performed differential gene expression analysis comparing aggregated RNA reads from each cell type against all others. Differential chromatin accessibility at marker genes showed broad concordance with expression differences (Figure 2D). Genomic tracks of the promoter regions of known marker genes demonstrated enriched chromatin accessibility alongside elevated RNA expression (Figure 2C), reinforcing assigned cell identities and validating the quality of the paired multi-omic data (Figure 2E).

### Cell composition changes in the human ASD putamen

We next analyzed cell-type composition (Figure S4G) to assess whether specific populations were altered in ASD/NDD samples relative to controls using Propeller^42^. The proportions of major cell populations, including medium spiny neurons, oligodendrocytes, and microglia, did not differ significantly between ASD/NDD and control samples (Figure 2F). In contrast, astrocytes were nominally significantly more abundant in ASD/NDD samples than controls (p = 0.014), which is consistent with the 16p11.2df mouse model (Figure 2F and Figure 1D). Subtype-level analysis revealed that this increase was primarily attributable to quiescent astrocytes (p = 0.0093), rather than reactive (*GFAP⁺*) astrocytes (p = 0.44) (Figure 2G). In addition, although the overall proportion of medium spiny neurons was unchanged, we detected small but nominally significant shifts in neuronal subtype composition: a cluster of D1 matrix neurons and a cluster of D1 striosome neurons were both modestly but significantly increased in autistic donors with a genetic diagnosis (p = 6.9 x10^-4^ and p = 0.047, respectively), with similar trends observed in autistic individuals without genetic diagnoses (Figure 2H).

### Cell type–specific transcriptional changes in human ASD putamen

To define transcriptional changes across putamen cell types in ASD/NDD, we performed differential gene expression analysis per cell type while controlling the effect of batch-related and other covariates using dreamlet^43^ (Methods). Our ASD/NDD cohort includes both cases with and without known genetic diagnoses, therefore, we compared each group to controls separately (Figure S5 A-B). We controlled the false discovery rate (FDR) at 5% (Benjamini– Hochberg), while also reporting nominally significant genes (p < 0.05) (Figure 3A, Figure S5C, Table S4). The most extensively differentially expressed genes were observed in donors with genetic diagnoses, consistent with the high effect sizes of such genetic variants in association with ASD/NDD (Figure 3A and Figure S5C). Among glia, astrocytes showed the greatest transcriptomic differences, with 666 DEGs passed FDR < 0.05. Within neurons, DEGs that met the FDR threshold were concentrated in D1 MSNs, rather than D2 MSNs, and were further enriched in the D1 striosome compartment at the subtype level (Figure 3A and Figure S5C). We identified 886 DEGs (317 upregulated, 569 downregulated) passed FDR < 0.05 in D1 striosome MSNs in donors with genetic diagnoses, and 320 DEGs (96 upregulated, 224 downregulated) in all ASD/NDD donors. Thus, ASD/NDD putamen samples had the largest transcriptional remodeling in D1 striosome MSNs. The compartment-specific dysregulation in striosomal D1 MSNs is notable given its role in integrating limbic input and gating dopaminergic reinforcement, aligning with evidence that aberrant striatal circuit function contributes to ASD-related behaviors^44,45^.

**Figure 3.**
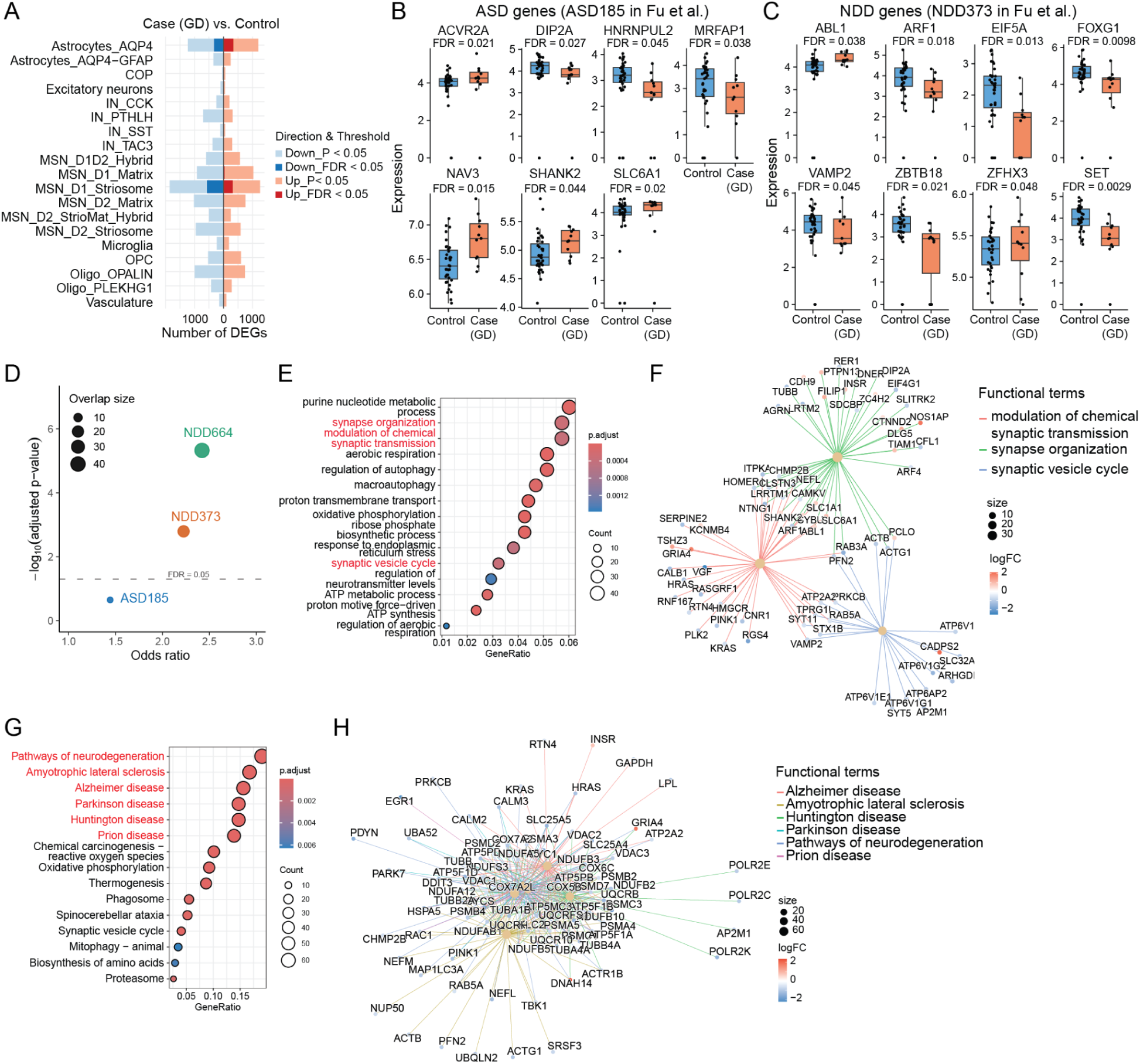
Cell type specific transcriptional alterations in ASD putamen. (A) Number and direction of DEGs in major cell types between autistic individuals with a genetic diagnosis and controls. (B) Box plots showing expression of DEGs of D1 striosome MSNs overlapping with ASD risk genes (ASD185 in Fu et al.). GD: Cases with genetic diagnosis. (C) Box plots showing expression of DEGs of D1 striosome MSNs overlapping with NDD risk genes (NDD373 in Fu et al.). Genes overlapped with ASD185 not shown. GD: Cases with genetic diagnosis. (D) Bubble plot showing enrichment of ASD and NDD gene sets with DEGs of D1 striosome MSNs. Odds ratio and p-value from Fisher’s exact test. P-value adjusted by BH method. Bubble size represents the number of overlapping genes between DEGs and each gene set. (E) GO biological process terms enriched for D1 striosome DEGs, including synaptic and metabolic processes. (F) Category–gene network showing the D1 striosome DEGs found in GO functional pathways. Dot size represents the number of DEGs found in the term. The color of nodes represents the fold change in cases with a genetic diagnosis versus controls. (G) KEGG enrichment of D1 striosome DEGs highlighting neurodegeneration pathways. (H) Category–gene network showing the D1 striosome DEGs found in KEGG functional pathways. Dot size represents the number of DEGs found in the term. The color of nodes represents the fold change in cases with a genetic diagnosis versus controls.

### Cell-type-resolved differential expression converges on synaptic programs of D1 striosome MSNs

For D1 striosome MSNs, we focused on the genes that are significantly altered in ASD/NDD donors with a genetic diagnosis (FDR < 0.05). We note that the ASD donors without a genetic diagnosis have a similar trend but lower magnitude (Figure S5D), suggesting a shared molecular signature that spans genetic architectures and diagnosis in this cell type. We intersected the DEGs from the D1 Striosome MSNs with a list of genes associated with ASD/NDD, identified in genome-wide mutation analyses by Fu et al. 2022^7^. Seven D1 striosome DEGs overlapped with high-confidence ASD risk genes (ASD185; Fu et al., 2022), including *ACVR2A*, *DIP2A*, *HNRNPUL2*, *MRFAP1*, *NAV3*, *SHANK2*, and *SLC6A1* (Figure 3B; Figure S6A). These genes are dysregulated, showing both higher or lower expression in cases with a genetic diagnosis as compared to controls, and are involved in a variety of biological processes (Figure 3B). *NAV3* encodes a neuron navigator protein implicated in axon guidance and neuronal migration; *ACVR2A* is a TGF-β receptor involved in BMP/activin signaling that can influence neuronal differentiation and synaptic plasticity; *SLC6A1* encodes a GABA transporter (GAT-1) responsible for reuptake of GABA from the synaptic cleft, and *SHANK2* encodes a postsynaptic density scaffold critical for excitatory synapse structure and function. We extended the intersection of DEGs to genes identified as a high risk in the context of NDDs (NDD373, NDD664 as defined by Fu et al.^7^) and discovered a significant overlap (Figure 3B-D, Figure S6 B-E). Consistent with the overall downregulation of genes in D1 MSNs, these genes were majorly downregulated in cases, 16/21 genes when compared to the NDD373 list, and 13/15 genes when compared with genes exclusive to the NDD664 list. Overall, these results identify key genes affected in the MSN D1 Striasome that are also known to have a genetic/mutational burden in NDD/ASDs.

Gene-ontology (Biological process) enrichment analyses of D1 striosome DEGs revealed significant enrichment of genes related to synapse organization (p.adjust = 3.2×10^-4^, gene ratio = 0.057), regulation of neurotransmitter levels (p.adjust = 1.5×10^-3^, gene ratio = 0.029), and the synaptic vesicle cycle (p.adjust = 3.6×10^-4^, gene ratio = 0.032), alongside oxidative phosphorylation (p.adjust = 1.33×10^-10^, gene ratio = 0.043), and ATP metabolic processes (p.adjust = 7.1×10^-5^, gene ratio = 0.028) (Figure 3E). This is consistent with the high energetic demands of MSN synapses and the coupling of neurotransmitter vesicle turnover to mitochondrial ATP production^45–47^. We analyzed synapse-related pathways using a category– gene network (Figure 3F), revealing that DEGs in D1 striosome MSNs converged on three coordinated synaptic programs, each showing a shared bias toward reduced synaptic efficacy: vesicle-cycle, chemical synaptic-transmission and synapse organization. Most genes within the vesicle cycle group, including *STX1B*, *SYT5*, *AP2M1*, *RAB3A*, *RAB5A*, *SLC32A1*, and multiple vacuolar ATPase subunits (*ATP6V1E1*, *ATP6V1G2*, *ATP6AP2*), were downregulated. These genes encode key components of synaptic vesicle docking, neurotransmitter loading, and recycling machinery, indicating a coordinated reduction in vesicle turnover and synaptic release capacity. The chemical synaptic-transmission term likewise trends downward across multiple signaling and modulatory genes involved in synaptic plasticity (*RASGRF1*, *HRAS*, *RGS4*, *CAMKV*, *HOMER1*, *PRKCB*), vesicle release and Ca²⁺ handling (*ATP2A2*, *VAMP2*, *CALB1*), and activity-dependent homeostasis (*PLK2*, *PINK1*, *RTN4*, *HMGCR*, *VGF*), indicating broad suppression of neurotransmitter release, postsynaptic signaling, and calcium-dependent synaptic plasticity within corticostriatal circuits. In contrast, the synapse-organization term shows mixed directionality. Core scaffolds and cytoskeletal regulators that maintain mature excitatory synapses (*HOMER1*, *CLSTN3*, *CAMKV*, *NEFL*, *CHMP2B*) and multiple adhesion or actin-associated molecules (*SLITRK2*, *DIP2A*, *AGRN*, *SDCBP*, *CFL1*) are downregulated, indicating weakened postsynaptic stability. In contrast, *SHANK2*, *SLC6A1*, and *CDH9* are upregulated together with remodeling and signaling factors (*TIAM1*, *DLG5*, *CTNND2*, *INSR*, *FILIP1*, *NOS1AP*), consistent with compensatory synaptic reorganization and cytoskeletal remodeling occurring alongside the loss of core scaffold strength. All three pathways converge on synaptic function, collectively suggesting reduced synaptic throughput and plasticity.

### D1 striosome MSNs in ASD with stress-vulnerable states analogous to early neurodegenerative conditions

Notably, KEGG enrichment analysis revealed that DEGs in D1 striosome MSNs were significantly enriched in multiple neurodegeneration-associated pathways, including Alzheimer’s disease (p.adjust = 3.5×10^-16^, gene ratio = 0.16), Parkinson’s disease (p.adjust = 2.0×10^-20^, gene ratio = 0.15), Huntington’s disease (p.adjust = 2.4×10^-18^, gene ratio = 0.15), prion disease (p.adjust = 2.9×10^-18^, gene ratio = 0.14), and amyotrophic lateral sclerosis (ALS) (p.adjust = 1.4×10^-19^, gene ratio = 0.17) (Figure. 3G–3H). These pathways converge on a shared network of mitochondrial oxidative phosphorylation, proteasomal subunits, and autophagy-related genes (e.g., *NDUFA5*, *ATP5F1D*, *COX6C*, *PSMA3*, *PINK1*, *PARK7*), the majority of which were downregulated in ASD. Downregulation of these genes indicates impaired mitochondrial and proteostatic functions and reduced energetic supply supporting neuronal integrity. This suggests that D1 striosome MSNs in ASD may exist in a low-energy, stress-vulnerable state analogous to early neurodegenerative conditions. Such a coordinated downregulation of energy metabolism and protein homeostasis likely compromises synaptic resilience and dopaminergic integration in these neurons, consistent with the energetic fragility of striatal circuits^45,46^ and prior evidence of mitochondrial suppression in ASD^48,49^. These analyses identify D1 striosome MSNs as a focal population with significant transcriptional changes across ASD etiologies, converging on synaptic pathways. This compartment-specific signature provides a mechanistic bridge between molecular pathology and basal-ganglia circuit dysfunction in ASD and nominates striosome D1 MSNs as a strategic target for circuit and therapeutic interrogation.

### Weakened synaptic support from putamen astrocytes in ASD

In homeostatic astrocytes (*AQP4*^+^, *GFAP*-Low), we observed 666 DEGs (338 upregulated, 328 downregulated) in donors with genetic diagnoses (Figure 4A). Notably, Astrocyte DEGs significantly intersected with all three sets of NDD/ASD genes as previously defined (Figure 3, Figure 4D). We identified 8 Astrocytic DEGs that overlapped with ASD185 (*ASXL3, H3F3A, MED13L, PACS1, SKI, SLC6A1, TNPO3, YWHAG*). These DEGs identified in astrocytes span multiple biological processes, including chromatin and transcriptional regulation (*ASXL3*, *H3F3A*, *MED13L*), protein trafficking and signaling (PACS1, YWHAG), TGF-β pathway modulation (*SKI*), GABA reuptake (*SLC6A1*), and nuclear import (*TNPO3*). Interestingly, *SLC6A1* is the only high-confidence ASD risk gene shared between astrocytes and D1 striosome MSNs, but it shows opposite directions of change, upregulated in MSNs and downregulated in astrocytes, highlighting cell type–specific regulation and functional divergence of the same ASD risk gene. In the intersection of astrocytic DEGs with NDD sets we found 30 DEGs overlapped with NDD373, and 38 DEGs overlapped with NDD664 sets, respectively. These DEGs were significantly altered in donors with a confirmed genetic diagnosis, and showed similar trends in ASD donors without a confirmed genetic diagnosis as well (Figure 4 B-C, Figure S7 A-B).

**Figure 4.**
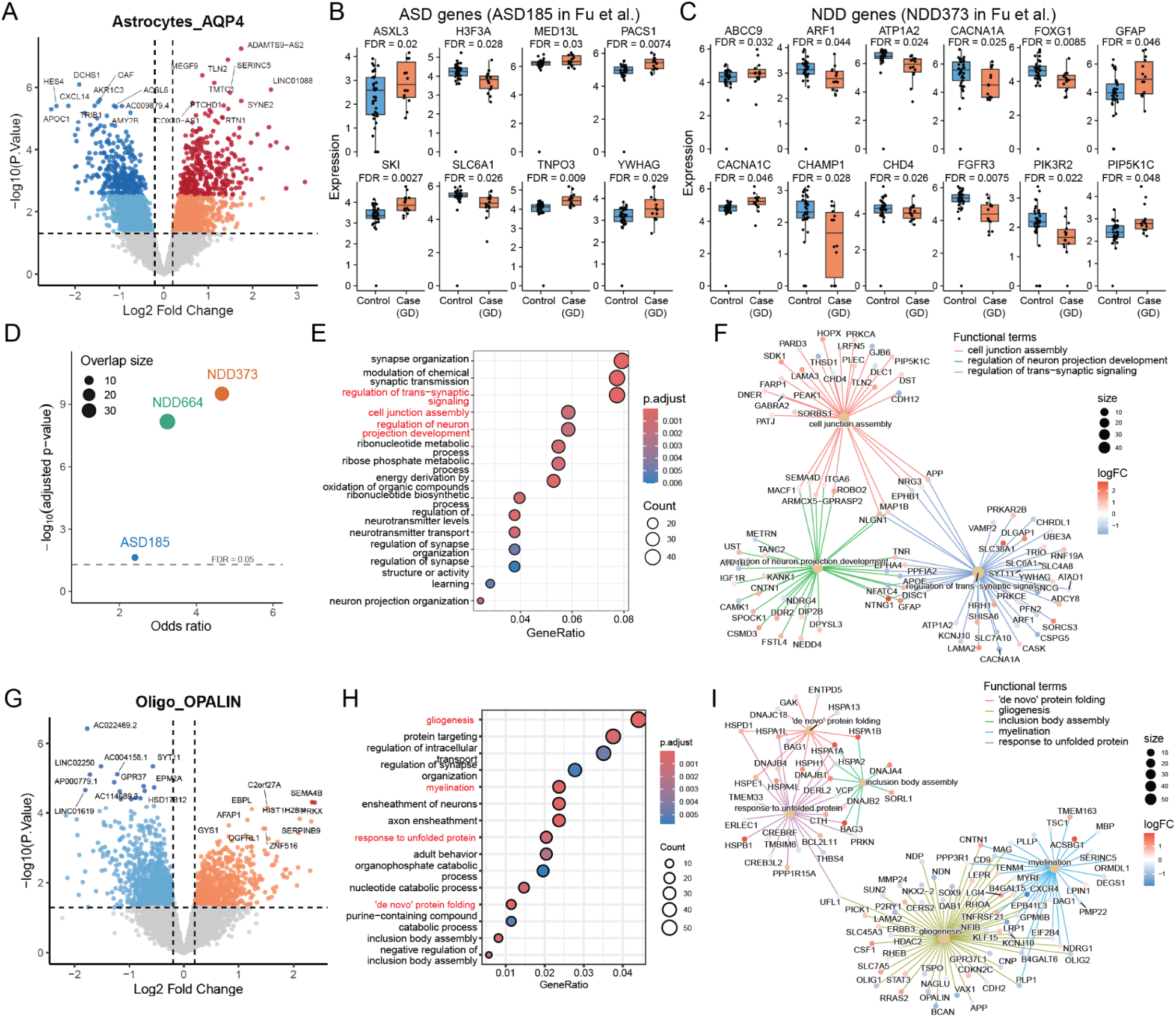
Glial transcriptional dysregulation in ASD putamen. (A) Volcano plot of DEGs in Astryctyes_*AQP4* showing upregulated and downregulated genes in ASD versus controls. Dark blue or red dots represent DEGs with FDR < 0.05. (B) Box plots showing expression of DEGs of Astryctyes_*AQP4* overlapping with ASD risk genes (ASD185 in Fu et al.). GD: Cases with genetic diagnosis. (C) Box plots showing expression of DEGs of Astryctyes_*AQP4* overlapping with NDD risk genes (NDD373 in Fu et al.). Genes overlapped with ASD185 not shown. GD: Cases with genetic diagnosis. (D) Bubble plot showing enrichment of ASD and NDD gene sets with DEGs of Astryctyes_*AQP4*. Odds ratio and p-value from Fisher’s exact test. P-value adjusted by BH method. Bubble size represents the number of overlapping genes between DEGs and each gene set. (E) GO biological process terms enriched for Astryctyes_*AQP4* DEGs, including trans-synaptic signaling and cell-junction assembly. (F) Category–gene network showing the Astryctyes_*AQP4* DEGs found in GO functional pathways. Dot size represents the number of DEGs found in the term. Color of nodes represents the fold change in cases with a genetic diagnosis versus controls. (G) Volcano plot of DEGs in *OPALIN*⁺ oligodendrocytes showing upregulated and downregulated genes in ASD versus controls. Dark blue or red dots represent DEGs with FDR < 0.05. (H) GO biological process terms enriched for nominal DEGs (P<0.05) of *OPALIN*⁺ oligodendrocytes, including gliogenesis and myelination. (I) Category–gene network showing the nominal DEGs (P<0.05) of *OPALIN*⁺ oligodendrocytes found in GO functional pathways. Dot size represents the number of DEGs found in the term. Color of nodes represents the fold change in cases with a genetic diagnosis versus controls.

Gene Ontology enrichment of DEGs in *AQP4*^+^ astrocytes mainly clusters into synapse-facing functions (Figure 4E): synapse organization, modulation of chemical synaptic transmission, regulation of trans-synaptic signaling, and cell-junction assembly; together with metabolic programs (ribonucleotide/ribose-phosphate metabolism, energy derivation/oxidation of organic compounds) and neurite support (neuron projection organization/development). Downregulated DEGs revealed a reorganization of astrocytic programs governing synaptic support and adhesion (Figure. 4F). Synaptic and signaling components, including *SLC6A1*, *VAMP2*, *CACNA1A*, *SYT11*, *ATP1A2*/*B1*, *KCNJ10*, *UBE3A*, *PPFIA2*, and *CAMK1*, were broadly downregulated, indicating reduced neurotransmission support and ionic homeostasis. In contrast, scaffolding and signaling regulators (*SLC38A1*, *DLGAP1*, *CASK*, *YWHAG*, *TRIO*, *HRH1*) and focal-adhesion and ECM-related genes (*TLN2*, *LAMA2*/*3*, *ITGA6*, *PLEC*, *DST*, *PIP5K1C*, *PRKCA*) were upregulated, reflecting a shift from broad synaptic support toward reinforced adhesive/ECM engagement consistent with reactive astrocyte remodeling and attenuated astrocyte–neuron communication.

KEGG enrichment of DEGs in *AQP4*⁺ astrocytes highlighted neurodegeneration-associated pathways (Alzheimer’s, Huntington’s, Parkinson’s, Prion diseases), core bioenergetic programs (oxidative phosphorylation, thermogenesis, carbon metabolism), and signaling/structural modules (cGMP-PKG, HIF-1, insulin signaling, GABAergic synapse, focal adhesion) (Figure S7C). Consistent with impaired metabolic support, the majority of respiratory-chain constituents were reduced, whereas adhesion and intracellular signaling terms displayed mixed directionality, suggesting compensatory remodeling rather than global activation. Mitochondrial/OXPHOS components (e.g., *NDUF*, *COX*, *ATP5* families, VDACs) form a downregulated hub that connects the disease pathways (Figure S7D). Energy- and stress-responsive programs (HIF-1, insulin signaling) bridge to this hub through glycolytic and hypoxia-sensing genes (e.g., *ALDOC*, *ENO1*, *PFKM*, *EGLN3*, *PDHA1*), while a partially segregated focal-adhesion cluster is anchored by ECM/adhesion regulators (e.g., *LAMA2/3/4*, *ITGA6*, *TLN2*, *SHC3*, *PIP5K1C*). Together, the topology indicates coordinated depression of astrocytic bioenergetics with concurrent adjustments in adhesion, an arrangement expected to diminish synaptic homeostasis and neuronal support.

### Disrupted myelination from putamen oligodendrocytes in ASD

In *OPALIN⁺* oligodendrocytes, our analysis revealed 17 DEGs that passed an FDR < 0.05 (Figure 4G). To evaluate the broad changes of oligodendrocytes in ASD/NDD, we included nominally significantly differentially expressed genes (p < 0.05; N=1733) in a gene-ontology enrichment analysis. GO enrichment of biological processes converges on two themes (Fig. 4H): myelin biology: gliogenesis, myelination/axon ensheathment, and regulation of synapse organization; and proteostasis/trafficking: protein targeting, regulation of intracellular transport, response to unfolded protein, *de novo* protein folding, and inclusion-body assembly. KEGG pathway analysis (Fig. S7 E-F) reinforces this architecture, highlighting protein processing in the endoplasmic reticulum. The category–gene network (Figure 4I) resolves these pathways into two interconnected modules: a myelination module anchored by structural and regulatory myelin genes (*MBP*, *PLP1*, *OPALIN*, *MAG*, *OLIG2, PMP22,* and *CNP*), and a proteostasis module centered on ER chaperones and co-chaperones (*HSPA1A/B/HSPH1*, *HSPA5*, *HSPD1*; DNAJ and BAG families), linked via trafficking components to protein-targeting terms and inclusion-body control (e.g., *ENTPD5*). Together, these results indicate that *OPALIN⁺* oligodendrocytes in ASD engage heightened ER/proteostatic and transport programs while remodeling adhesion and lipid metabolism, reflecting a stress-compensatory rather than myelin-generative state. This transcriptomic profile suggests that oligodendrocytes may be attempting to preserve and repair existing myelin sheaths under cellular stress, consistent with defective or inefficient myelin maintenance^50^.

### Linking Differential Gene Expression to Cell Type-Specific Cis-Regulatory Landscapes

To mechanistically explain transcriptional changes, we investigated upstream regulatory elements and regulatory factors using snATAC-seq data. First we performed peak calling separately for each cell type and generated a unified consensus peak set with 1,166,673 peaks (see methods). Across all cell types, we observed an extensive set of common as well as distinct cell type–restricted peaks (Figure 5A), with D1 and D2 medium spiny neurons (MSNs) contributing the largest numbers of unique peaks (n=181,696). Genomic annotation revealed an enhancer-dominant chromatin landscape: most peaks resided within intronic or distal intergenic regions, whereas less were located near promoters (Figure 5B). Analysing the difference of genomic distribution of peaks called from neuronal cells or non-neuronal cells, we found the peaks called from non-neuronal cells have higher ratio from promoter (22.1%) compared with neuronal cells (18.73%).

**Figure 5.**
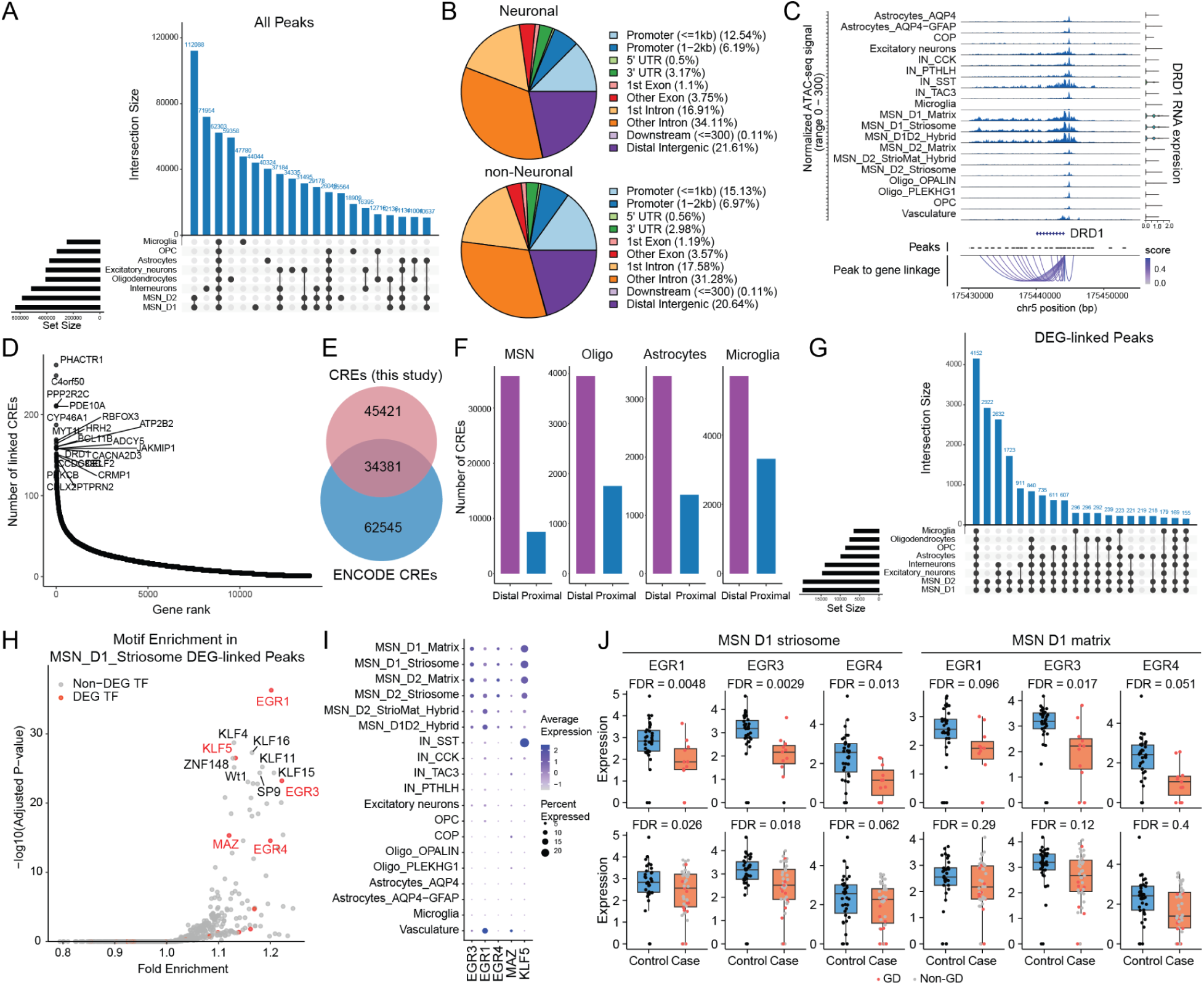
Linking transcriptional alterations to cis-regulatory elements and transcription factor programs in ASD putamen. (A) Intersection of ATAC-seq peaks across major putamen cell types showing both shared and cell type–specific accessibility profiles. (B) Distribution of genomic locations peaks called from neuronal population or non-Neuronal populations. (C) Example locus (*DRD1*) showing peak to gene linkage inferred from the correlation of accessibility of regulatory elements with gene expression. (D) Ranking of genes by the number of linked cis-regulatory elements (CREs), showing a long-tailed distribution. (E) Vennplot showing the number of shared CREs between this study and the published ENCODE dataset. (F) Comparison of distal versus proximal CREs across cell-type specific peaks. Cell type– specific peaks were identified based on chromatin accessibility relative to all other cell types, with thresholds of log fold change > 2 and adjusted p < 0.05. (G) Intersection of DEG-linked peaks across cell types. (H) Motif enrichment in CREs associated with DEGs in D1 striosome MSNs.TFs within the DEG of ASD vs. controls in D1 striosome MSN are shown are highlighted. (I) Dot plot showing expression of TFs enriched in DEG-associated peaks across cell types. Dot size represents the percentage of nuclei expressing each gene, and color intensity denotes scaled average expression. (J) Box plots showing expression of *EGR3* and *EGR1* in D1 striosome MSNs and D1 matrix MSNs. GD: Cases with genetic diagnosis. Non-GD: Cases without genetic diagnosis.

To identify candidate cis-regulatory elements (CREs) for each gene, we inferred peak–gene associations by correlating chromatin accessibility with gene expression across matched cell populations (Methods). At the DRD1 locus, multiple distal elements were co-accessible with a single gene, illustrating the characteristic many-to-one enhancer architecture of neuronal transcriptional regulation (Figure 5C). Ranking genes by the number of linked CREs revealed a long-tailed distribution, with neuronal and synaptic genes (as *PPP2R2C, PDE10A, CYP46A1, MYT1L, HRH2, BCL11B, ADCY5, RBFOX3, ATP2B2, AKAP5, DRD1*) exhibiting the densest enhancer neighborhoods (Figure 5D), consistent with the complex regulatory architecture required for activity-dependent transcription in the striatum, which is a specific feature for neurons^51^. To evaluate the CREs we identified, we compared it with another peakset of human putamen published on ENCODE from DNase-seq^52^, and found 43% peaks overlapped (Figure 5E).

To characterize the distribution of CREs across cell types, we identified cell type–specific CREs by comparing their chromatin accessibility with that of all other cell types. We then assessed the genomic distribution of these CREs, distinguishing between proximal (<2 kb from the TSS) and distal regions linked to their target genes (Figure 5F). Interestingly, CREs associated with MSNs showed a markedly higher contribution from distal elements compared with those linked to glial cells such as oligodendrocytes, astrocytes, and microglia. This suggests that transcriptional regulation in MSNs relies more heavily on distal regulatory elements, whereas glial gene regulation is more biased toward proximal promoter regions, which has similar trends in other brain regions^53^.

With established CRE–gene linkages, we identified those CREs associated with DEGs. DEG-associated peaks were shared across multiple cell types, with the majority found in MSNs (Figure 5G). This ratio of shared regulatory elements suggests that common enhancer circuits, rather than strictly cell type–specific elements, may underlie convergent transcriptional dysregulation in ASD/NDD, reflecting cross-cell-type coordination in striatal gene regulation.

We next considered which transcription factor (TF) programs are encoded within DEG-associated accessible regions. Motif enrichment analysis of DEG-linked peaks in D1 striosome MSNs (using peaks linked to expressed genes in the same population as background) revealed strong enrichment for early growth response (EGR) family members (*EGR1*, *EGR3*, *EGR4*), together with KLF family zinc-finger regulators (*KLF4*, *KLF5*, *KLF16*) and additional transcription factors such as *MAZ* and *SP9* (Figure 5H). Among these, *EGR1, EGR3* and *EGR4* were all significantly downregulated in D1 striosome MSNs in ASD donors, with a similar downward trend in D1 matrix MSNs (Figure 5J), implicating them as central regulators of transcriptional programs disrupted in ASD. Both TFs were selectively expressed in MSNs (Figure 5I), consistent with their roles as immediate-early genes coupling synaptic activity to downstream transcriptional responses. This suggests a broad attenuation of EGR-mediated enhancer activity and reduced transcriptional responsiveness in the striatum in ASD/NDD.

### ASD-associated regulatory programs driven by EGR3 and EGR1 in D1 striosome MSNs

To further delineate the regulatory architecture underlying transcriptional differences in ASD putamen, we leveraged our single-nuclei multiome dataset to infer transcription factor–target gene networks using SCENIC+^54^, which integrates motif presence within cis-regulatory elements and the co-variation between transcription factor expression, chromatin accessibility, and target gene transcription. From these data, we constructed enhancer-based regulons (eRegulons) and quantified their activity across cell types using both target-gene and target-region signatures (Methods). The target-gene activity reflects the expression of all genes assigned to a given TF regulon, and the target-region score reflects the accessibility of regulatory elements linked to those genes. Each eRegulon was classified as activating or repressing according to the direction of TF–target associations (Figure 6A–6B).

**Figure 6.**
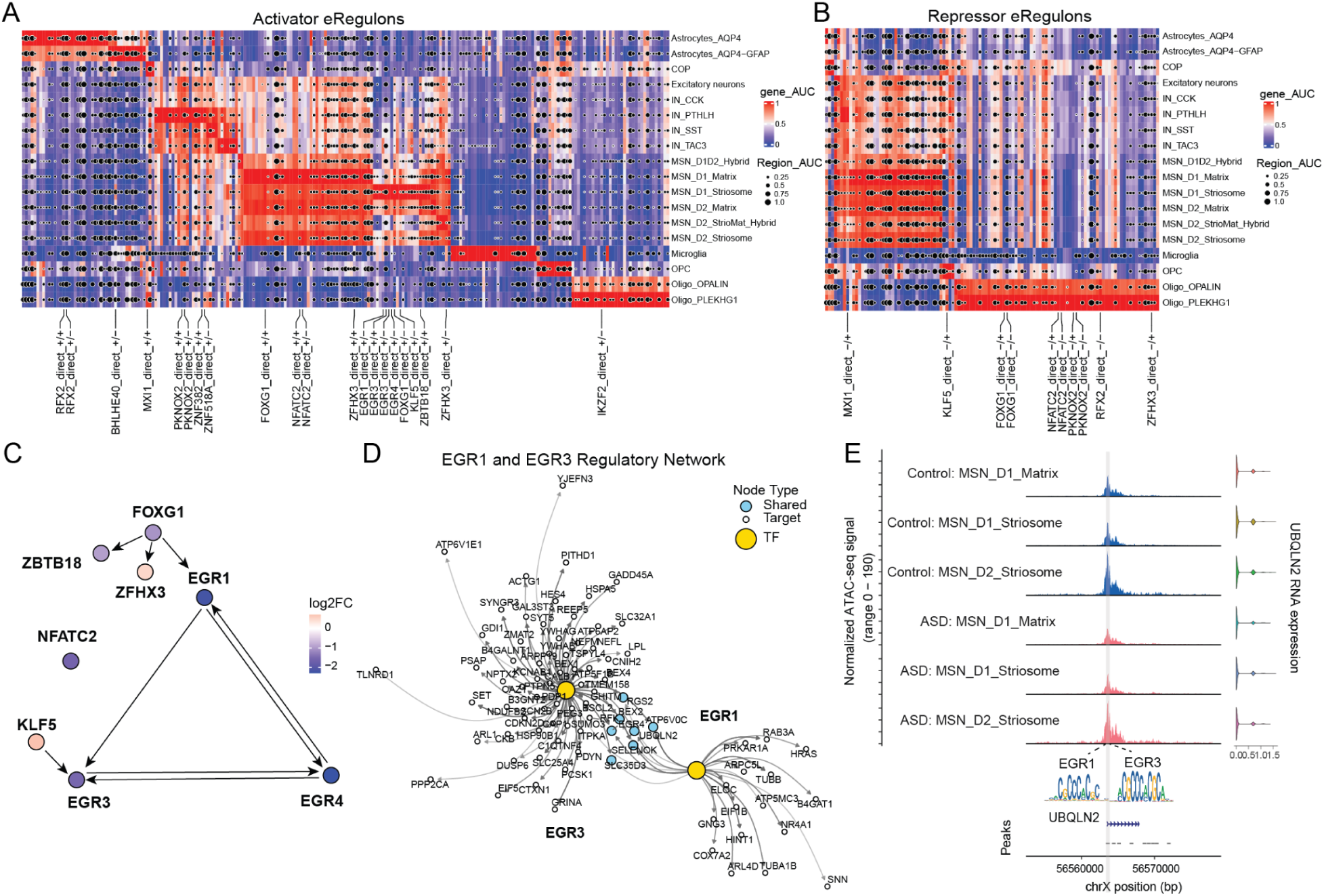
*EGR3* and *EGR1*-centered regulatory programs in ASD putamen. (A) Heatmap showing the gene-based AUC scores and region-based AUC scores of activator eRegulons across cell types. Color represents gene-based AUC scores. Dot size represents region-based AUC scores. TFs within the DEG of ASD vs. controls in D1 striosome MSN are shown are highlighted. (B) Heatmap showing the gene-based AUC scores and region-based AUC scores of repressor eRegulons across cell types. Color represents gene-based AUC scores. Dot size represents region-based AUC scores. TFs within the DEG of ASD vs. controls in D1 striosome MSN are shown are highlighted. (C) Regulatory network among TFs with D1 striosome MSN specific eRegulons. TFs within the DEG of ASD vs. controls in D1 striosome MSN are shown. Color of nodes represents the fold change in cases with a genetic diagnosis versus controls. (D) Regulatory network of *EGR1* and *EGR3*. Color of nodes represents different types of nodes. (E) A representative locus (*UBQLN2*) of shared targets of *EGR1* and *EGR3*. The regulatory region with *EGR1* and *EGR3* motifs is highlighted.

We next focused on TFs showing differential expression in D1 striosome MSNs of ASD donors. Among the activator eRegulons, we identified a transcriptional module comprising multiple ASD/NDD-dysregulated transcription factors, including immediate early and activity-dependent regulators (*EGR1*, *EGR3*, *EGR4*, *KLF5*, and *NFATC2*), together with additional zinc-finger and neuronal regulators (*ZBTB18*, *FOXG1*, and *ZFHX3*) (Figure 6A). This module showed its highest activity in MSNs, particularly within the D1 striosome subtype, aligning with the locus of strongest transcriptional alteration in ASD. Among these factors, *EGR3* and *EGR1* stood out as the most prominent: both were previously identified in motif enrichment of DEG-associated peaks (Figure 5H) and were significantly downregulated in MSNs in ASD/NDD (Figure 5J).

*EGR3* (Early Growth Response 3) is a neuronal activity–dependent transcription factor that acts downstream of calcium and BDNF signaling to control synaptic plasticity and experience-driven gene expression. It belongs to the immediate-early EGR family with *EGR1*, *EGR2*, and *EGR4*, which are rapidly induced by neuronal activity and dopamine signaling. Among these, protein coding mutations in *EGR3* have been identified among ASD individuals^55,56^. Their reduced activity suggests weakened coupling between neuronal activity and gene-regulatory responses in ASD.

By extracting the regulatory network among DEG transcription factors whose eRegulon activities are highly specific in MSNs, we found that *EGR3*, *EGR4*, and *EGR1* form a mutually connected regulatory circuit (Figure 6C). Upstream of this EGR circuit, *FOXG1* emerged as a key regulatory node that also controls *ZFXH3* and *ZBTB18*. Notably, *FOXG1*, *ZFXH3*, and *ZBTB18* are well-established neurodevelopmental disorder genes included in the curated NDD373 list from Fu et al 2022^7^.

To map their downstream regulatory landscape, we reconstructed the *EGR3*/*EGR1*-centered gene regulatory network (Figure 6D). The resulting network connected *EGR3* and *EGR1* to targets involved in synaptic organization (e.g., *GRINA*, *SYNGR3, PSAP*), axon guidance and cytoskeletal remodeling (e.g., *RGS2*, *TUBB*, *ARPC5L*), and metabolic or mitochondrial regulation (e.g., *ATP6V1E1*, *COX7A2*). These targets collectively converge on neuronal activity–dependent and energy-coupled transcriptional programs critical for maintaining MSN excitability and synaptic homeostasis (Figure S8).

Notably, in the downstream of shared targets of *EGR3* and *EGR1* (Figure 6D), *UBQLN2* is also significantly differentially expressed (logFC = -1.02, FDR = 0.02) in D1 striosome MSNs, which is reported to be critical in neurodegeneration diseases^57,58^. Inspection of representative loci of *UBQLN2* demonstrated reduced chromatin accessibility, coinciding with the predicted *EGR3*/*EGR1* binding motifs at regulatory elements (Figure 6E). *UBQLN2* is a ubiquitin-binding shuttle protein that serves as a molecular hub connecting synaptic maintenance and cellular energy homeostasis. It regulates proteostasis at synapses and supports mitochondrial quality control, thereby coordinating protein turnover and energy balance essential for neuronal function^57,58^.

This finding supports a model in which activity-induced *EGR3* and *EGR1* act as transcriptional activators linking neuronal firing to synaptic and metabolic gene expression. In ASD/NDD, the downregulation of *EGR3*/*EGR1* together with accessibility at their enhancer targets likely impair this coupling, leading to attenuated synaptic responsiveness and disrupted homeostatic regulation in the striatum. These results place *EGR3* and *EGR1* at the center of a convergent regulatory cascade that integrates activity, metabolism, and transcription, providing a mechanistic bridge between altered neuronal activity and the molecular pathology of ASD/NDD in the putamen.

### Cell-type-specific cis-regulation at ASD risk loci

To investigate how ASD-associated genetic risk variants may generate cell-type-specific regulatory effects, we integrated ASD GWAS summary statistics^7,59–83^ with our single-nucleus ATAC-seq data. Linkage disequilibrium score regression (LDSC) revealed significant enrichment (p = 0.0038) of ASD heritability within open chromatin regions of MSNs, indicating that these neuronal populations are key substrates of ASD genetic liability (Figure 7A).

**Figure 7.**
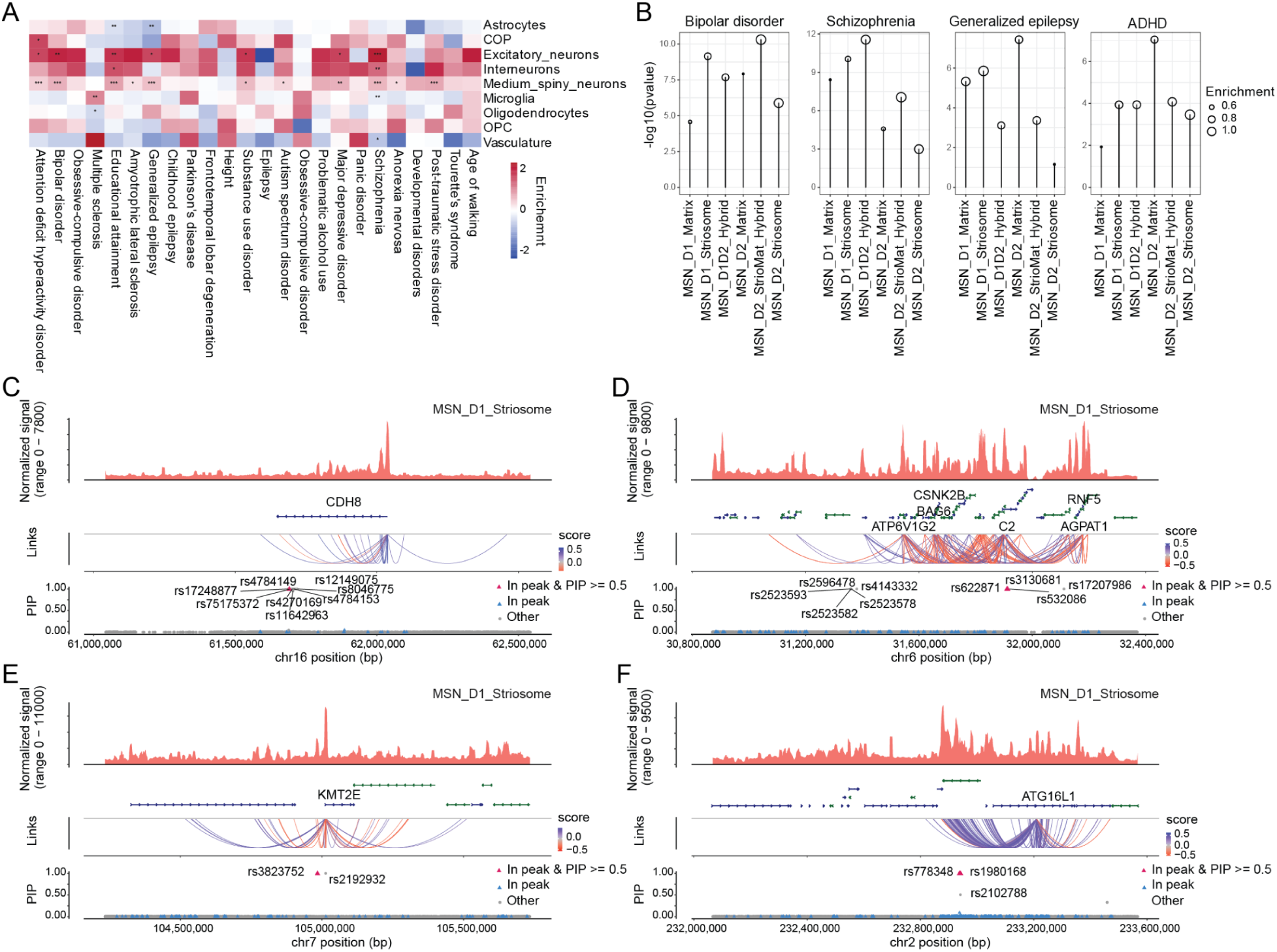
Cell-type-specific cis-regulation at ASD-related disease genetic risk loci. (A) LDSC enrichment of various neuropsychiatric GWAS traits, including ASD, in open chromatin across cell types(* FDR < 0.05, ** FDR < 0.01, *** FDR < 0.001). (B) Lollipop plot showing LDSC enrichment of Bipolar, Schizophrenia, Generalized Epilepsy and ADHD GWAS among subtypes of MSN. (C-F) Cis-regulatory architecture at the following GWAS loci in MSN D1 Striosome: lead ADHD SNP (rs16963957) and lead SCZ SNPs (rs1078323, rs2857693, rs3823752). Genes correlated with CREs involving causal SNPs are highlighted. Only MSN D1 Striosome CREs were used to fine-map SNPs for ADHD. For the fine-mapping track, we only labeled SNPs with PIP > 0.5. The SNP located within a gene-correlated MSN D1 Striosome CRE and exhibiting a high PIP value is marked with a red triangle.

Notably, medium spiny neurons also displayed strong heritability enrichment across multiple psychiatric disorders, including schizophrenia (SCZ) (p = 4e-7), bipolar disorder(p = 1e-6), generalized epilepsy (p = 1e-5) and attention-deficit/hyperactivity disorder (ADHD) (p = 1e-5), highlighting shared neuronal regulatory mechanisms across neurodevelopmental conditions^6,84,85^ (Figure 7A). These disorders are known to share genetic risk overlap and convergent neurobiological features with ASD, suggesting common molecular and regulatory vulnerabilities. Additionally, different subtypes of MSN, including D1 striosome MSNs, showed significant and selective enrichment patterns specific to individual psychiatric disorders (Figure 7B).

We next performed fine-mapping of ASD-, ADHD-, and SCZ-associated single-nucleotide polymorphisms (SNPs) within cell type–specific CREs of D1 striosome MSNs to estimate their posterior inclusion probabilities (PIPs). Interestingly, variants near *CDH8* exhibited strong posterior inclusion probabilities of being causal (PIP > 0.5) and overlapped gene correlated CREs (r = 0.38), suggesting direct transcriptional regulation of their target genes (Figure 7C). Notably, *CDH8*-associated variants were fine-mapped from ADHD loci but overlapped neuronal CREs enriched for ASD-related MSN D1 striosome. *CDH8* is one of the most established genes associated with ASD risk that encodes a synaptic adhesion molecule^86–87^, suggesting that ADHD-linked regulatory variation at this locus may converge on molecular pathways relevant to ASD pathophysiology. Additionally, several SCZ fine-mapped causal SNPs (rs11208758, rs622871, rs3130681, and rs3823752; PIP > 0.5) were identified within CREs of D1 striosome MSNs. These CREs are found to be linked with ASD-differentially expressed genes, including *C2*, *BAG6*, *ATP6V1G2*, *RNF5*, *AGPAT1*, and *ATG16L1* (Figure 7 D-F). Notably, *KMT2E* and *CSNK2B* (FDR = 0.06 in the differential gene expression test), both previously reported NDD-related genes from the NDD373 list, were also linked to peaks containing causal SNPs from SCZ. Together, this integrative SNP-CRE-gene framework, these loci point to convergent regulatory mechanisms across psychiatric disorders, implicating striatal MSNs, particularly D1 striosome subtypes, as a key cell population mediating shared genetic risk among SCZ, ADHD, and ASD.

## Discussion

In this study, we applied an unbiased approach for lineage tracing to compare neurodevelopmental phenotypes across multiple brain regions in the developing brain. Leveraging a well-characterized genetic mouse model of ASD, we show widespread perturbations of development trajectories across the brain, but most notably within the striatum, a brain structure involved in motor control, learning and emotion. Our results are consistent with prior observations of increased number of MSNs in 16p11.2 heterozygous deletion^88^, and moreover our clonal barcoding analysis suggests that the expansion of MSNs originates from progenitors forming regionally dispersed clones, which also produce neurons migrating toward the OB. Altered abundance of D1 matrix and striosome MSNs was also detected in a subset of individuals diagnosed with ASD compared to individuals without ASD diagnosis. This altered abundance could reflect differences in early development consistent with findings in the mouse model.

D1 striosome MSNs exhibited the strongest transcriptional changes in ASD/NDD. Altered genes were enriched in synaptic vesicle cycling, neurotransmission, and postsynaptic scaffolding, core processes for corticostriatal plasticity and reinforcement learning. These findings align with the known role of striosome circuits in integrating limbic input and modulating dopaminergic reward signaling, linking subcortical dysfunction to ASD-related behavioral inflexibility. Notably, striosome organization and expression of dopamine receptor 1 are significantly altered between mice and humans^89^, and striosome MSNs may play an outsized role in limbic control of motor behaviors. Pathway analysis of gene expression changes within D1 striosome MSNs revealed downregulated mitochondrial and proteostatic programs, indicative of a reduced energetic capacity and stress resilience. These findings suggest that D1 striosome MSNs neurons operate in a low-energy, vulnerable state analogous to early neurodegenerative conditions, which may be a consequence in ASD. This altered state of vulnerability might be consistent with recent findings of increased incidence of neurodegenerative disorders among autistic individuals^90,91^.

By integrating epigenomic and transcriptomic signatures, we demonstrate that differential expression in ASD is primarily driven by enhancer modulation. Transcription factor binding motif enrichment and regulatory inference analyses highlighted downregulation of gene regulatory programs regulated by early growth response factors *EGR3* and *EGR1* as central transcriptional regulators of genes involved in synaptic organization, cytoskeletal remodeling, and mitochondrial function. These findings position *EGR3* and *EGR1* as pivotal regulators that integrate neuronal activity, transcriptional adaptation, and metabolic homeostasis in the striatum. Disruption of this axis likely weakens dopaminergic feedback and corticostriatal plasticity, offering a unified framework linking molecular, cellular, and circuit-level dysfunction in ASD/NDD. The disrupted program was most pronounced in D1 striosome MSNs, likely because these neurons uniquely regulate dopaminergic tone via direct projections to the substantia nigra pars compacta. Their role in reward evaluation and action selection makes them particularly vulnerable to disruptions in activity-dependent transcriptional programs such as the EGR pathway, offering a mechanistic basis for striosome-selective vulnerability in ASD. EGR3 has been independently linked to ASD through rare de novo gene disrupting mutations^55,56^.

Although similar disruption to *EGR3* mediated networks did not emerge in our analysis of 16p11.2 mice, it is possible that such effects could emerge later in development, or be specific to humans. Prior studies of *Egr3* in mice have implicated this gene in mediating gene expression regulation of NMDA receptor signaling^92^, and overexpression of this gene in striatal MSNs has been implicated in striatal behaviors^93^.

*EGR3* and *EGR1*, members of the immediate-early growth response transcription factor family, are rapidly induced in striatal MSNs following dopaminergic signaling through D1-type receptors, where they couple neuronal activity to long-term transcriptional and synaptic adaptations. *EGR3*, in particular, mediates dopamine- and calcium-dependent transcriptional programs that regulate MSN excitability, dendritic spine remodeling, and reward-related learning^94^. Loss or dysregulation of *Egr3* impairs synaptic plasticity and produces behavioral abnormalities overlapping with ASD-related phenotypes^92,95^. *EGR3* has been also reported as a master regulator of genes differentially expressed in the brains of patients with neuropsychiatric illnesses ranging from schizophrenia and bipolar disorder to Alzheimer’s disease^96–98^. *EGR1* has been reported to regulate key synaptic genes such as *SHANK3*^99^ and is necessary for dopaminergic signaling during social behavior^100^. These *EGR* transcription factors form a dopamine-responsive regulatory hub in MSNs that coordinates synaptic and transcriptional plasticity, providing a mechanistic link between striatal dysfunction and ASD-associated alterations in neural circuit regulation.

Glial populations showed coordinated remodeling that complements neuronal stress. Astrocytes shifted from synaptic and metabolic support toward adhesion and extracellular-matrix engagement, indicating diminished capacity for neurotransmission maintenance. Oligodendrocytes exhibited altered myelination, proteostasis, and intracellular transport consistent with a compensatory, stress-responsive state. Notably, astrocyte abundance was consistently increased in both the 16p11.2df mouse model and the human putamen. Together, these results suggest a disturbance in cellular cooperation within the striatum. Altered gene expression, abundance, and neurophysiology of astrocytes has been previously observed in postmortem tissue^101^, as well as several models of ASD including valproic acid^102^. Moreover, astrocytes are known to express several important ASD-linked molecular programs implicated in support of synaptic transmission^103,104^.

In addition, our findings also point to a cross-disorder convergence at both genetic and molecular levels. Our functional analysis of DEGs revealed that ASD shares core molecular dysfunctions with other psychiatric and neurodegenerative disorders such as Alzheimer, Parkinson, Huntington, and ALS. Moreover, GWAS SNP analysis demonstrated significant enrichment of genetic risk factors associated with ASD, ADHD, SCZ, bipolar disorder, and epilepsy in the CREs of MSN, underscoring the shared genetic and molecular architecture among these conditions. These convergent patterns suggest that ASD and other psychiatric disorders may involve common regulatory pathways and cellular stress responses, implying that overlapping therapeutic strategies could address similar neurobiological phenotypes across diagnoses.

Together, our study reveals that developmental lineage dynamics, transcriptional dysregulation, and enhancer remodeling converge to disrupt striatal homeostasis in ASD/NDD. The multi-omic resource established here broadens ASD/NDD research beyond its traditional cortical focus, emphasizing subcortical regulatory mechanisms as key contributors to the disorder. Our findings position *EGR1* and *EGR3* as central regulators that couple neuronal activity to synaptic and metabolic gene programs, linking dopaminergic signaling to striatal function. Future work should dissect how EGR-mediated transcriptional networks are dynamically tuned during development. This work provides a framework for developing targeted interventions to stabilize striatal circuitry and improve functional outcomes in ASD/NDD.

## Acknowledgments

We are grateful to all members of the Nowakowski lab for their insight and advice while conducting these experiments and preparing the manuscript. We would like to thank the UCSF Helen Diller Family Comprehensive Cancer Center Flow and Cell Sorting Core Facility (RRID:SCR_026372) for their assistance with FACS. We would also like to thank Ryan Delgado for his pioneering work in the initial design of STICR.

We would like to thank Alan Packer and Marta Benedetti of the Simons foundation, Kelly Gleason (UT Southwestern), David Amaral and team at UC Davis and Autism Brain Net for their efforts. https://www.autismbrainnet.org/tissue-application/. Tissues and data were obtained from Autism BrainNet, a resource of the Simons Foundation Autism Research Initiative (SFARI). Autism BrainNet also manages the Autism Tissue Program (ATP) collection, previously funded by Autism Speaks. We are grateful and indebted to the families who donated tissue for research purposes to Autism BrainNet and the ATP.

Multiome libraries were sequenced at the UCSF Center for Advanced Technology (CAT), supported by UCSF PBBR, RRP IMIA, and NIH 1S10OD028511-01 grants. This project was supported by the National Institute of Mental Health (NIMH) and National Institute of Neurological Disorders and Stroke (NINDS) with grant numbers U01MH130962, R01NS123263 and R01MH128364, the National Institutes of Health (NIH) with grant numbers U01MH122681, R01MH125516, and R01MH129751, the Health Data Research UK QQ2 Molecules to Health Records Driver Program, the Simons Foundation Autism Research Initiative Sex Differences Collaboration (SFARI #736613), the California Institute for Regenerative Medicine (CIRM) DISC0-14429, as well as gifts from Esther A. & Joseph Klingenstein Fund, the Shurl and Kay Curci Foundation, the Sontag Foundation, William K. Bowes Jr. Foundation.

T.J.N. is a New York Stem Cell Foundation Robertson Neuroscience Investigator.

This publication was supported by and coordinated through the Brain Initiative Cell Atlas Network (BICAN).

## Author contributions

Conceptualization: G.Y., V.S., E.M.W., S.J.S., T.J.N.

Data curation: E.M.W., M.G., G.Y., V.S., Y.H., N.R., J.W., D.S.P., F.L., N.S., S.D.

Formal analysis: G.Y., E.M.W., Y.H., V.S., M.G.

Funding acquisition: T.J.N., S.J.S.

Investigation: G.Y., V.S., R.L., M.R.S., C.E.

Methodology: G.Y., V.S., E.M.W., R.L., M.R.S., C.E.

Project administration: G.Y., V.S., E.M.W., Y.H.

Resources: T.J.N., S.J.S.

Software: Y.H.

Supervision: T.J.N., S.J.S.

Visualization: G.Y., Y.H., E.M.W., V.S.

Writing – original draft: G.Y., V.S., E.M.W., Y.H.

Writing – review & editing: G.Y., V.S., E.M.W., Y.H., M.R.S., M.G., N.R., D.S.P., S.J.S., T.J.N.

## Competing interests

Y.H. is a co-founder and equity holder of Neptune Bio. All other authors declare no competing interests.

## SUPPLEMENTARY FIGURES

**Figure S1.**
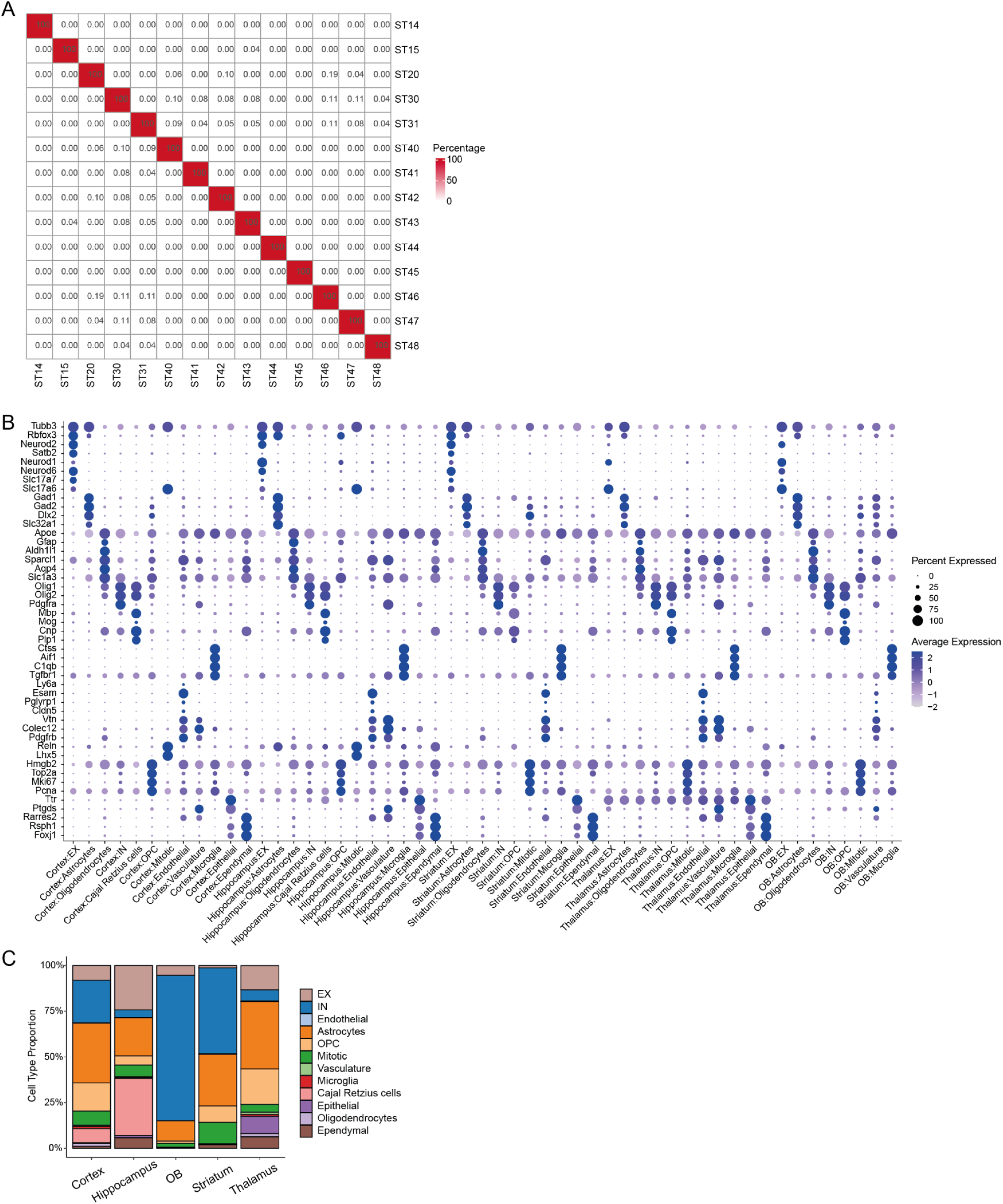
Characterization of Virus-Labeled Cells with Lineage Barcodes, related to Figure 1. (A) Heatmap showing the percentage of identical clones shared between pairs of brains. For each comparison, the percentage was calculated relative to the sample with the lower total clone count. Each row or column indicates an independent brain. (B) Dot plot showing the expression of marker genes of different cell types across regions. (C) Stacked bar plot showing the proportion of cell types in each brain region.

**Figure S2.**
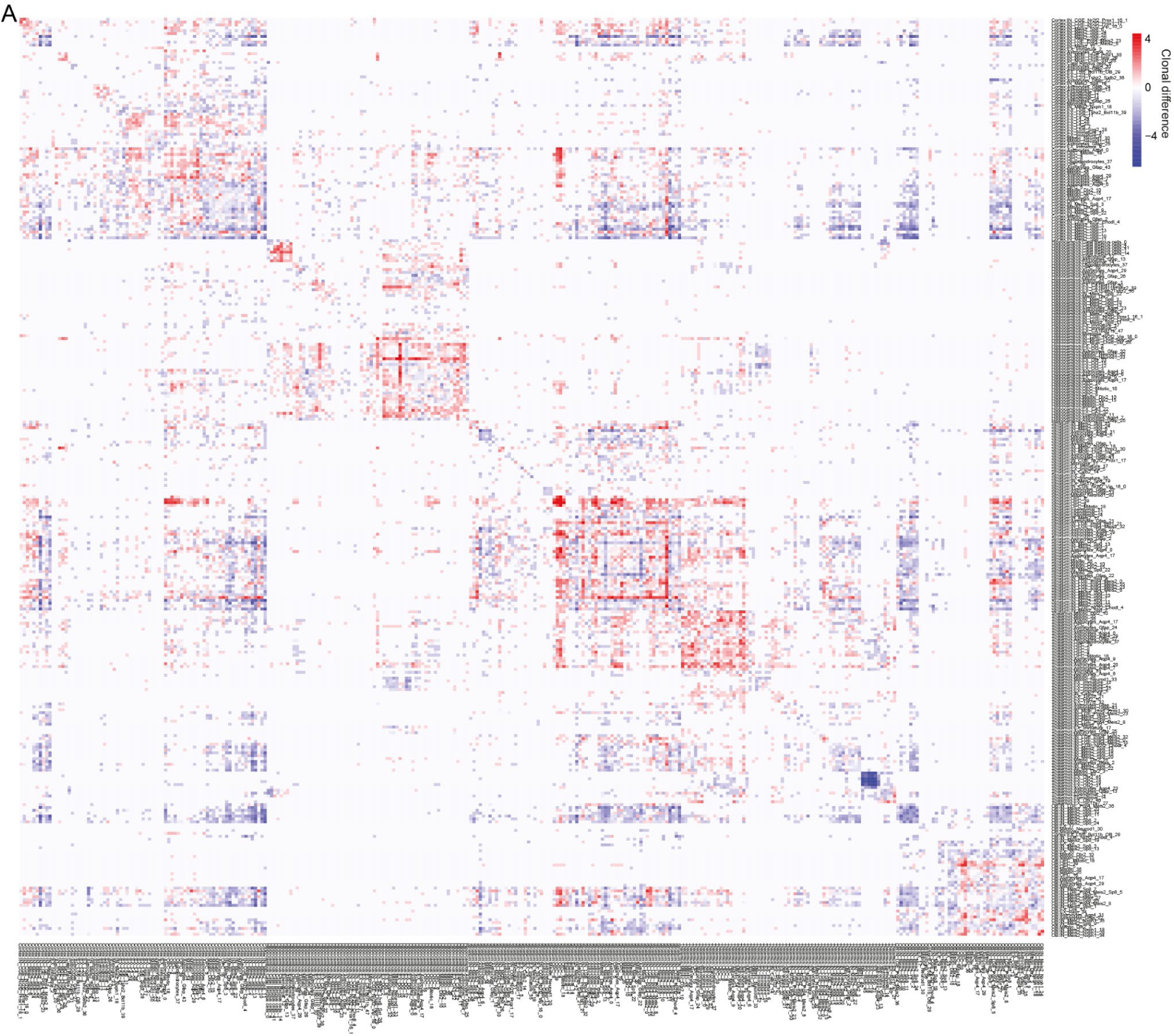
Differential clonal coupling between 16p11.2df and wild-type mice, related to Figure 1. (A) Heatmap showing the difference in clonal coupling of cell clusters across all regions between 16p11.2df and wild-type mice injected at E12.5. Colors represent changes in clonal coupling, calculated as the difference between normalized shared clonal counts. All clusters are shown.

**Figure S3.**
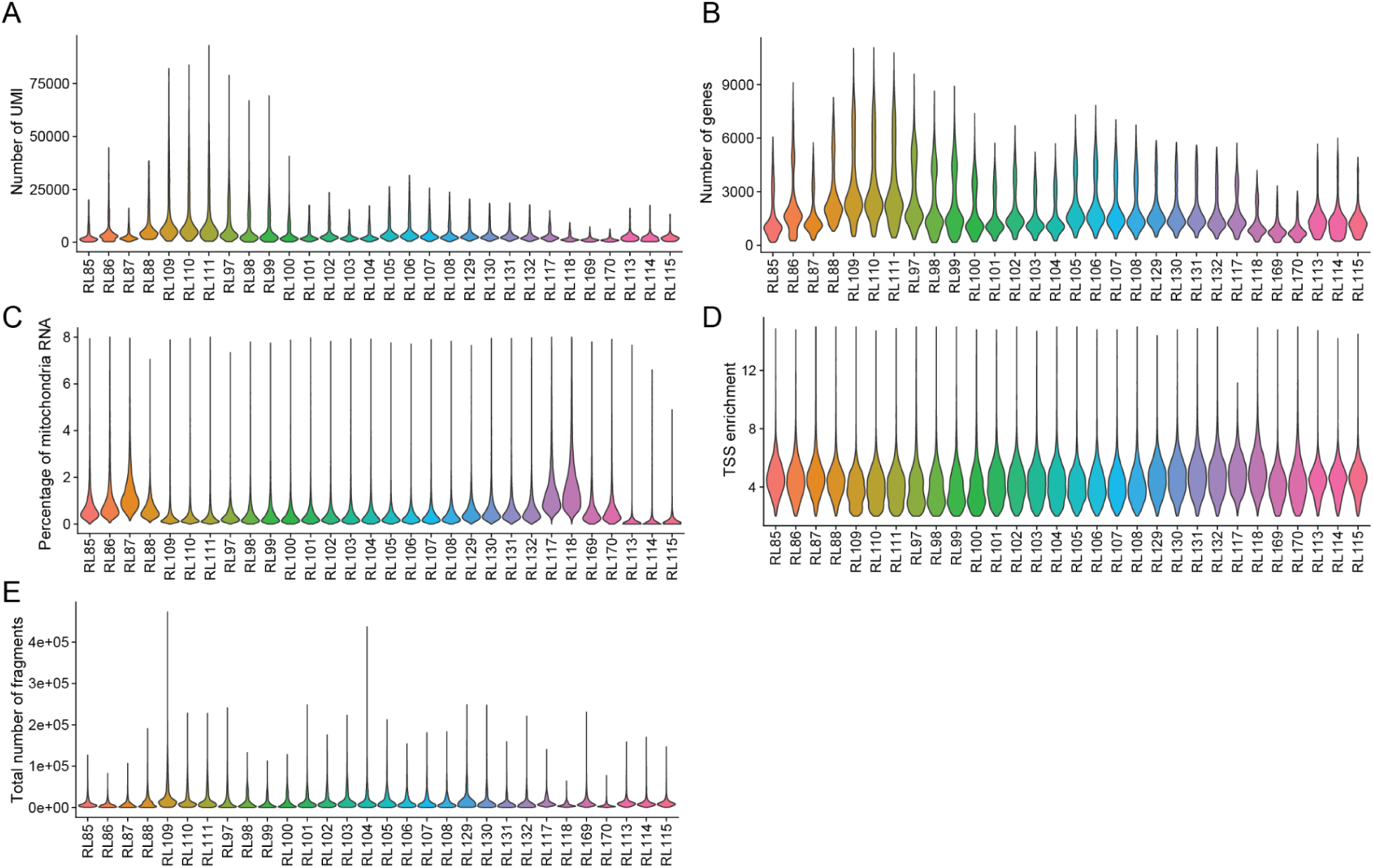
Quality control of snMultiome data of human putamen, related to Figure 2. (A) Violin plot showing the number of UMIs detected per cell in each well of the 10x experiments. (B) Violin plot showing the number of genes detected per cell in each well of the 10x experiments. (C) Violin plot showing the percentage of mitochondrial RNA per cell in each well of the 10x experiments. (D) Violin plot showing TSS enrichment scores of ATAC-seq data per cell in each well of the 10x experiments. (E) Violin plot showing the number of ATAC-seq fragments captured per cell in each well of the 10x experiments.

**Figure S4.**
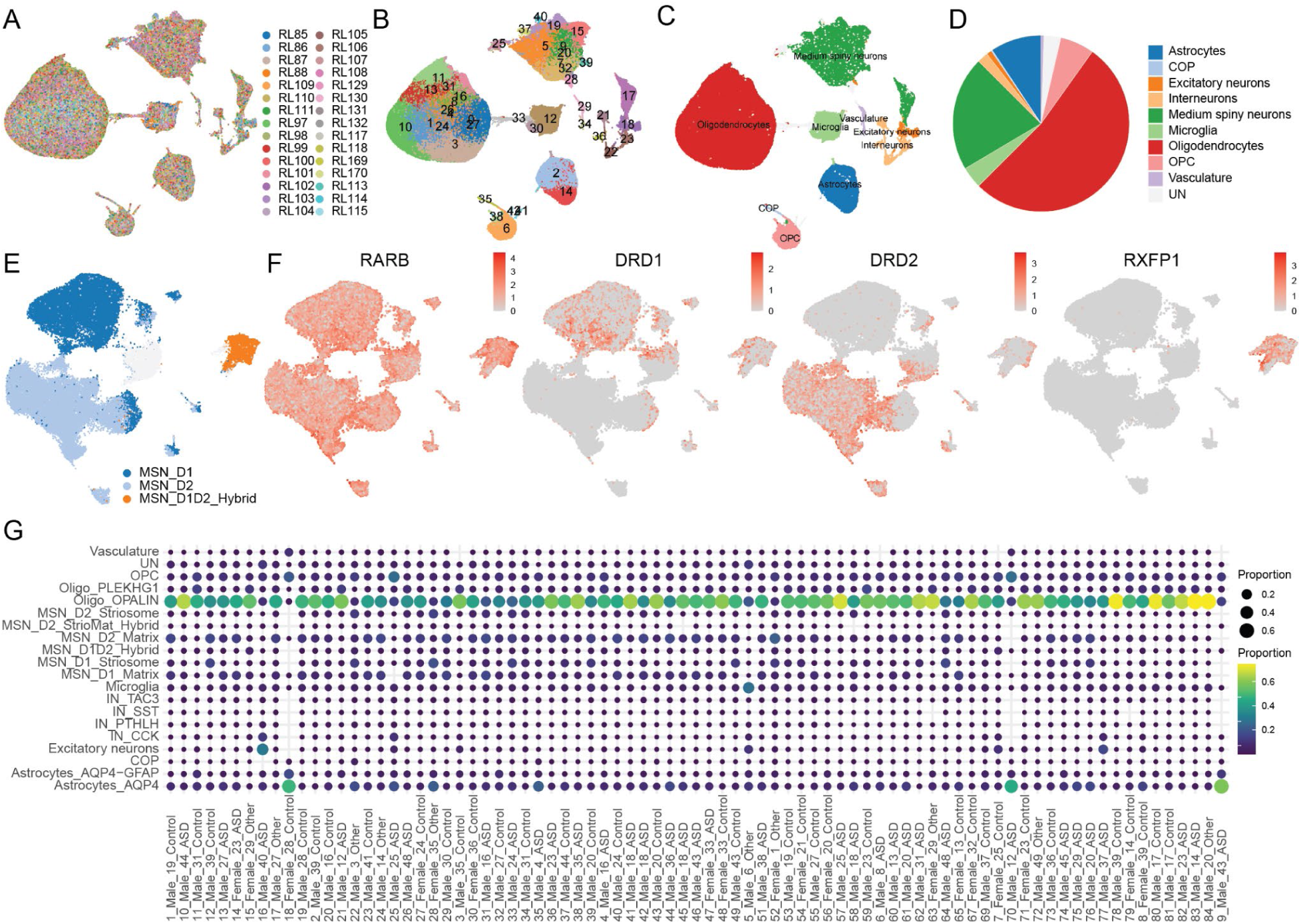
Characterization of Cell Types in Human Putamen Data, related to Figure 2. (A) UMAP visualization of high-quality nuclei colored by wells from the 10x Genomics experiments. (B) UMAP visualization of high-quality nuclei colored by clusters. (C) UMAP visualization of high-quality nuclei colored by annotated cell types. (D) Bar plot showing the proportion of each cell type. (E) UMAP visualization of medium spiny neurons (MSNs), with colors indicating MSN subtypes. (F) UMAP visualization showing the expression of marker genes distinguishing MSN subtypes. (G) Dot plot showing the proportion of each cell type across individual donors.

**Figure S5.**
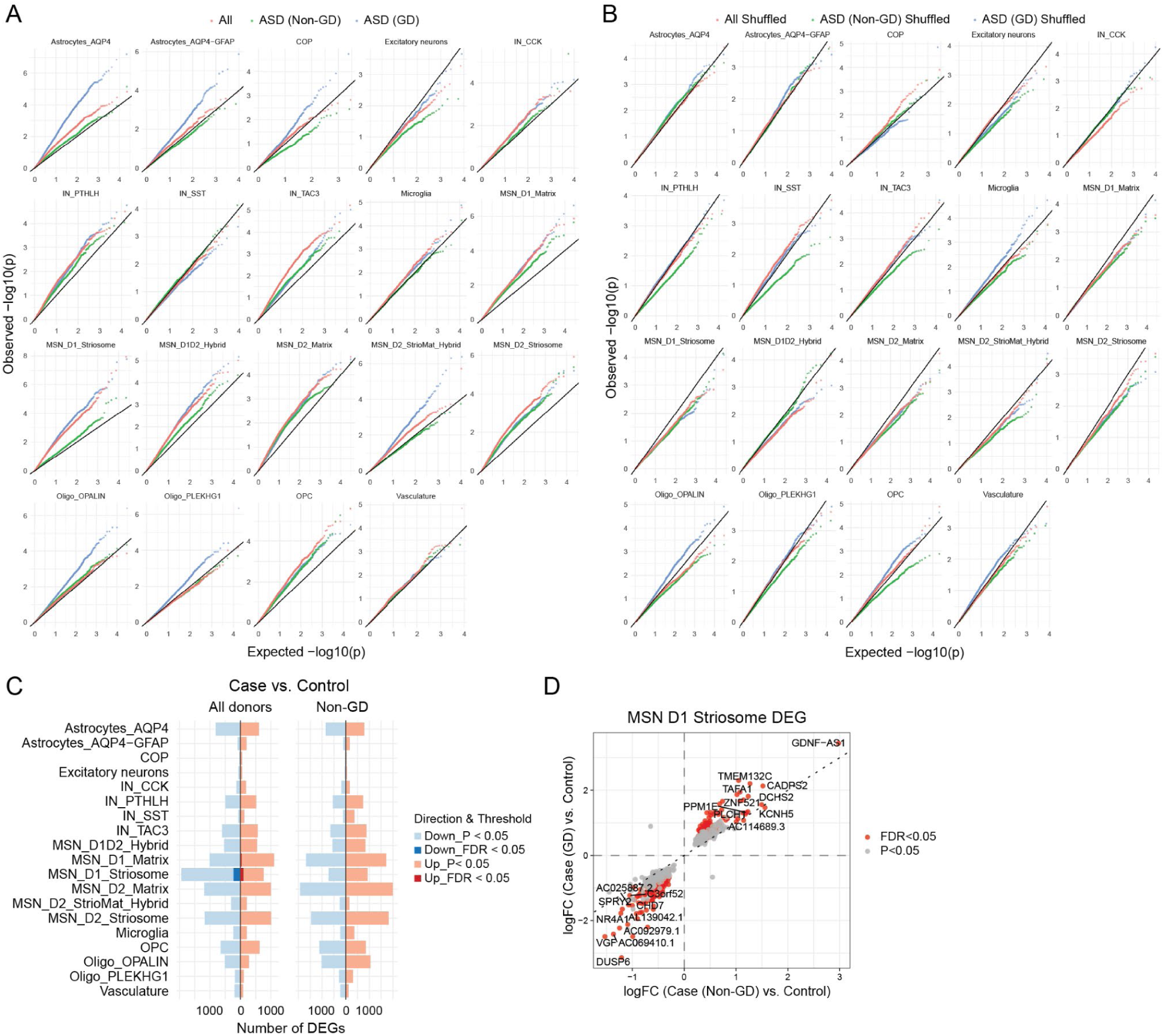
Differentially expressed genes in different cell types, related to Figure 3. (A) Quantile-quantile (QQ) plots showing the distribution of p-values from differential gene expression analyses comparing all ASD cases vs. controls, ASD (non-genetic diagnosis) vs. controls, and ASD (genetic diagnosis) vs. controls, using real sample labels. (B) QQ plots showing the corresponding distributions after random shuffling of sample labels, serving as a null control for comparison. The x-axis represents the expected −log10(p) values under the null hypothesis, and the y-axis represents the observed −log10(p) values from the differential expression tests. Deviations above the diagonal line indicate enrichment of significant genes beyond what is expected by chance. (C) Numbers and directions of differentially expressed genes (DEGs) in major cell types comparing ASD (All, Non-GD) versus controls. (D) Concordant DEGs between GD and Non-GD ASD in D1 striosome MSNs.

**Figure S6.**
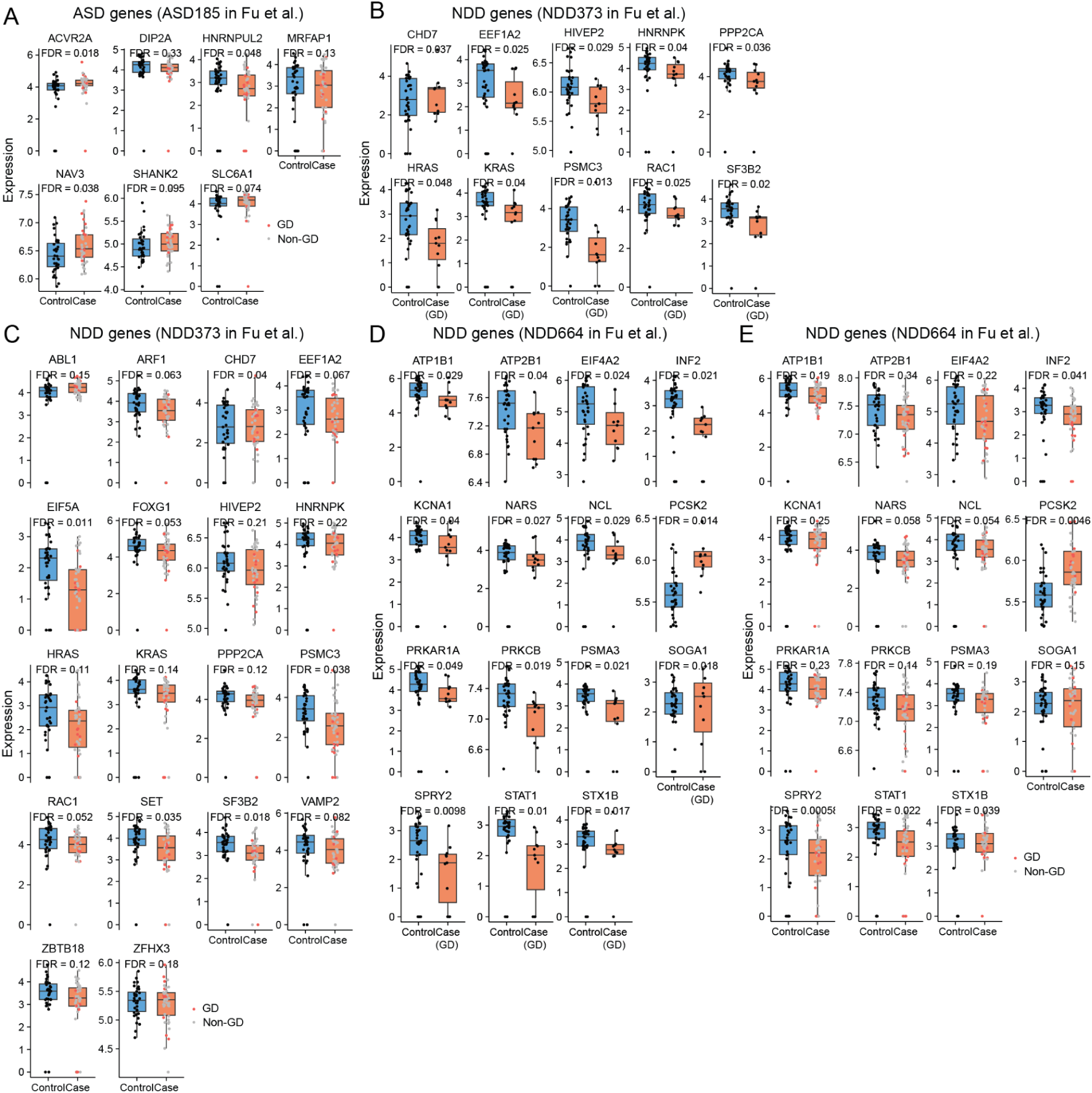
Expression of DEG of D1 striosome MSNs overlapping with ASD/NDD risk genes, related to Figure 3. (A) Box plots showing expression of DEGs of D1 striosome MSNs overlapping with ASD risk genes (ASD185 in Fu et al.). GD: Cases with genetic diagnosis. Non-GD: Cases without a genetic diagnosis. (B) Box plots showing expression of DEGs of D1 striosome MSNs overlapping with NDD risk genes (NDD373 in Fu et al.). Genes overlapped with ASD185 not shown. GD: Cases with genetic diagnosis. (C) Box plots showing expression of DEGs of D1 striosome MSNs overlapping with NDD risk genes (NDD373 in Fu et al.). Genes overlapped with ASD185 not shown. GD: Cases with genetic diagnosis. Non-GD: Cases without a genetic diagnosis. (D) Box plots showing expression of DEGs of D1 striosome MSNs overlapping with NDD risk genes (NDD664 in Fu et al.). Genes overlapped with ASD185 and NDD373 not shown. GD: Cases with genetic diagnosis. (E) Box plots showing expression of DEGs of D1 striosome MSNs overlapping with NDD risk genes (NDD664 in Fu et al.). Genes overlapped with ASD185 and NDD373 not shown. GD: Cases with genetic diagnosis. Non-GD: Cases without a genetic diagnosis.

**Figure S7.**
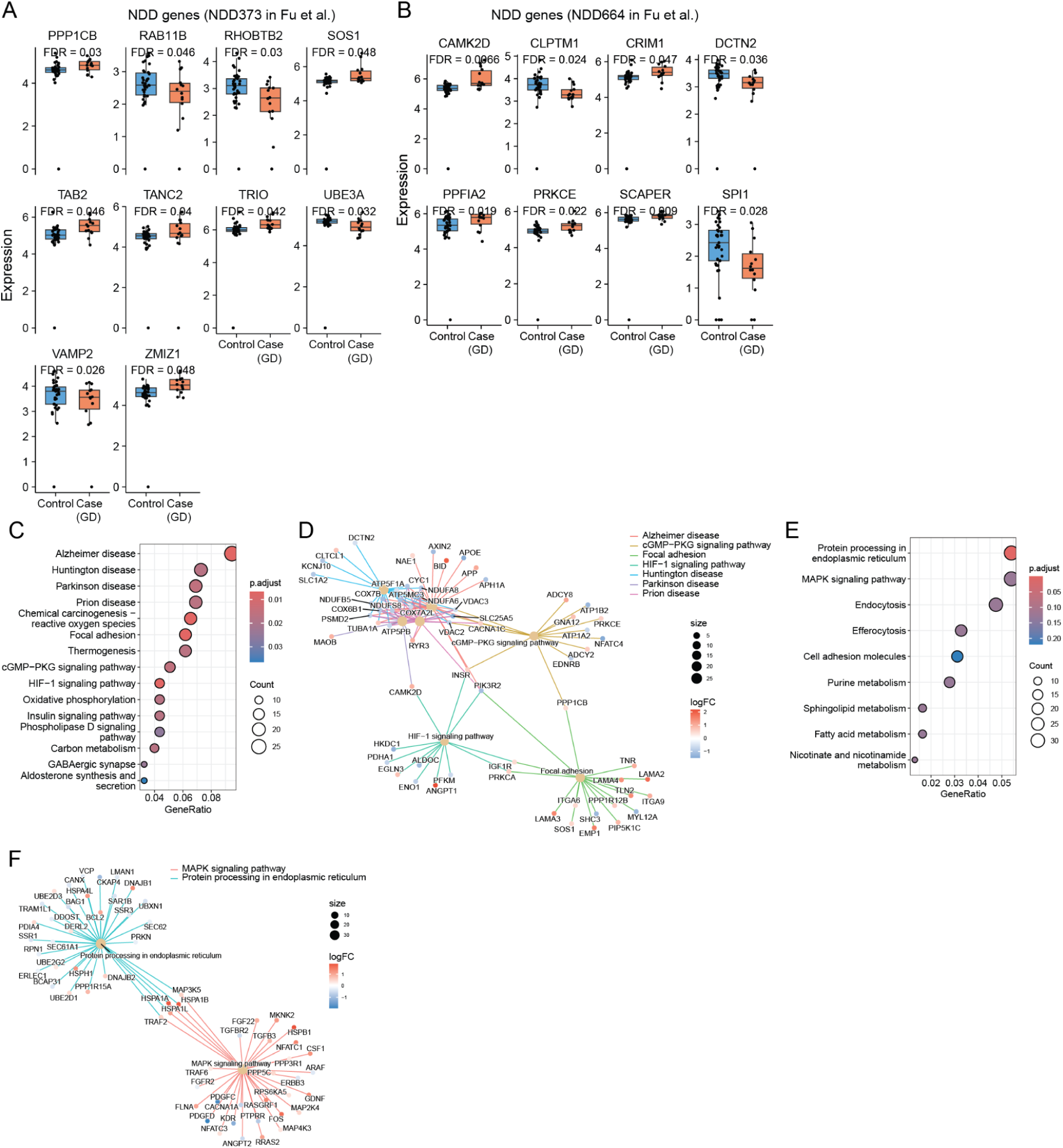
Differentially expressed genes in glia, related to Figure 4. (A) Box plots showing expression of DEGs of Astrocytes_*AQP4* overlapping with ASD risk genes (NDD373 in Fu et al.). Genes overlapped with ASD185 not shown. (B) Box plots showing expression of DEGs of Astrocytes_*AQP4* overlapping with NDD risk genes (NDD664 in Fu et al.). Genes overlapped with ASD185 and NDD373 not shown. (C) KEGG pathways enriched for Astryctyes_*AQP4* DEGs. (D) Category–gene network showing the Astryctyes_*AQP4* DEGs found in KEGG pathways. Dot size represents the number of DEGs found in the term. Color of nodes represents the fold change in cases with a genetic diagnosis versus controls. (E) KEGG pathways enriched for nominal DEGs (P<0.05) of *OPALIN*⁺ oligodendrocytes, (F) Category–gene network showing the nominal DEGs (P<0.05) of *OPALIN*⁺ oligodendrocytes found in KEGG pathways. Dot size represents the number of nominal DEGs found in the term. Color of nodes represents the fold change in cases with a genetic diagnosis versus controls.

**Figure S8.**
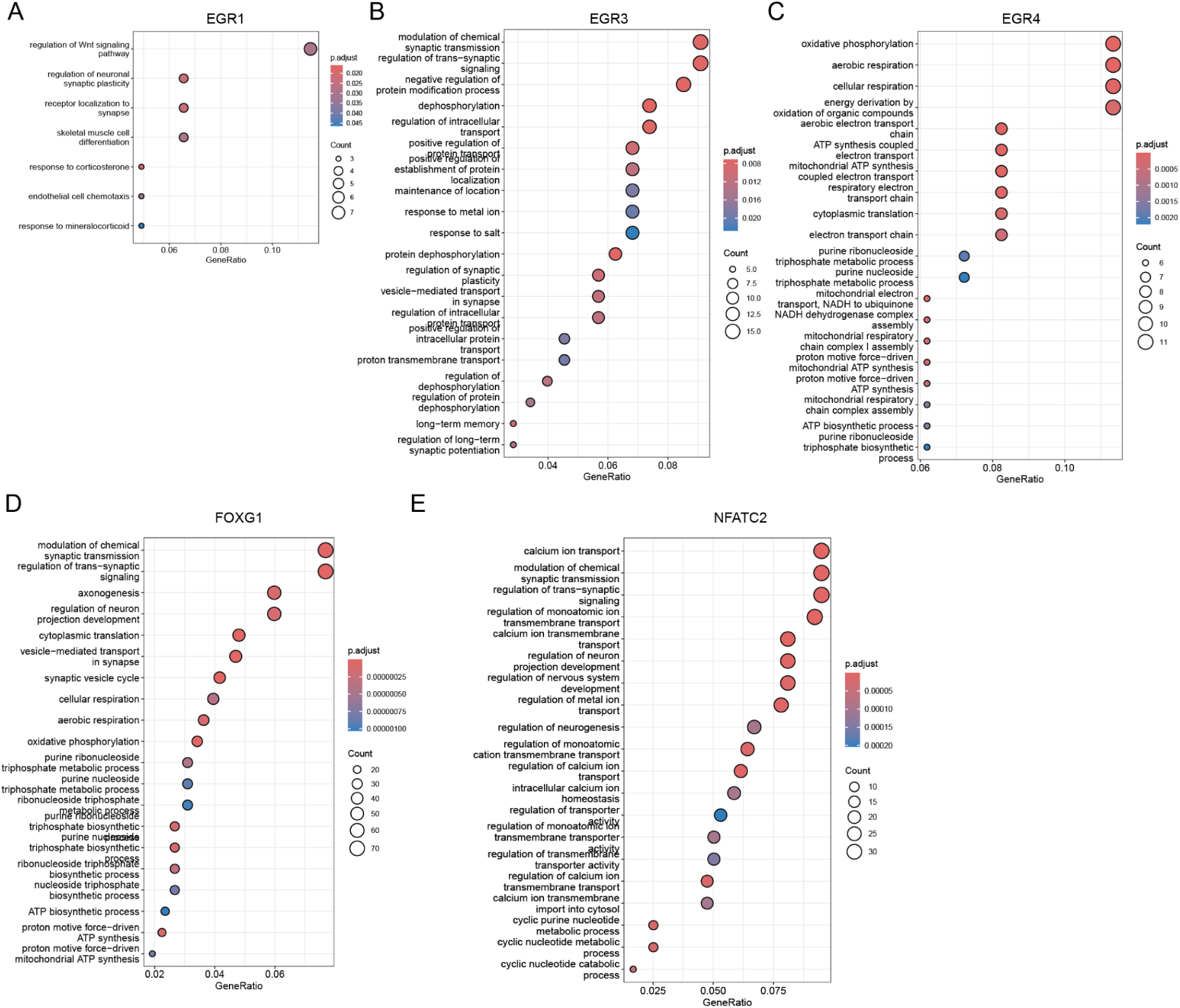
Function enrichment of regulatory target genes of key TFs, related to Figure 6. GO biological process terms enriched in the regulatory target genes of (A) *EGR1*, (B) *EGR3*, (C) *EGR4*, (D) *FOXG1*, (E) *NFATC2*.

## METHODS

### Plasmid and Lentivirus production for STICR library

STICR barcode plasmid library was prepared as previously described(addgene, #180483, #186334, #186335). STICR lentivirus was produced using a third-generation lentivirus packaging system: pMDLg/pRRE (Addgene, catalogue number 12251), pRSV-Rev (Addgene, 12253) and VSVG envelope (Addgene, 12259). Plasmids were transfected into Lenti-X HEK293T cells (Takara Bio, 632180) using Lipofectamine 3000 (Thermo Fisher, L3000075). In order to improve the viral titre, we also co-transfected pcDNA3.1 puro Nodamura B2 plasmid (Addgene, 17228) along with the other plasmids. Lenti-X 293T cells were grown and transfected in Dulbecco’s modified Eagle’s medium (DMEM) (Fisher, MT10017CV) supplemented with 10% fetal bovine serum (FBS) (Hyclone, SH30071.03) and 1% penicillin streptomycin (Fisher, 15070063). Twelve to eighteen hours after transfection, media was replaced with DMEM supplemented with 2% FBS (Hyclone, SH30071.03), 1% penicillin streptomycin, and 1x ViralBoost Reagent (Alstem, VB100). Forty-eight hours after transfection, media were collected, passed through a 0.45 μM filter (Corning, 431220), and then ultracentrifuged at 22,000g for 2 h. Pellets were resuspended in 100 μl of sterile phosphate-buffered saline (PBS) (Thermo, 14190250) overnight at 4 °C and then aliquoted and stored at −80 °C.

### Mice and in utero surgeries

All animal procedures were approved by the Institutional Animal Care and Use Committee (IACUC). C57BL/6 wild-type females were purchased from Charles River Laboratories(California, United States). 16p11.2df (B6129S-Del(7Slx1b-Sept1)4Aam/J, #013128) mice and B6129SF1/J (#101043) mice were purchased from The Jackson Laboratory. 16p11.2df mice were bred to B6129SF1/J hybrid mice each generation. All animals are maintained at the University of California, San Francisco. For timed pregnant mice, the plug date was designated as E0.5. In utero surgery and injection of the STICR lentiviral library in the lateral ventricles of the embryonic mouse forebrains were performed as previously described51. Pregnant dams were kept in single cages, and pups before P21 were kept with their mothers until collection.

### Tissue collection of the mouse brain

Virally injected brains were collected from mouse pups at postnatal day 4. Brains were dissected in ice-cold pre-bubbled artificial cerebrospinal fluid (aCSF) and sectioned into coronal sections. Each brain region was manually dissected out from sliced brains based on the guidance of the anatomy atlas from the Allen Institute. Collected tissue was then dissociated in the solution of papain (Worthington, LK003178) supplemented with 0.5 mg/ml DNase (Sigma Aldrich, #DN25-1G) diluted with HBSS (Gibco,14025134) at 37 °C. The tissues were incubated at 37 °C for 30-60 min, with gentle trituration every 15 min until samples had no visible chunks. Papain was quenched by adding 1%BSA (Militeny) to the solution. Samples were spun down and resuspended in HBSS + 0.1% BSA and passed through the 35 μm filter into a FACS tube (Falcon # 352235). To isolate and collect virally infected cells, flow cytometry was done using a BD FACSAria III Cell Sorter (BD FACSDiva Software, version 8.0.2) with a 100-µm nozzle. The cell suspensions were first gated on forward scatter, then within this population based on eGFP expression. eGFP-expressing cells were sorted into HBSS + 0.1% BSA for downstream processing on the 10x Genomics Chromium platform. Samples were kept on ice before and after FACS, and the chamber and collection tubes were maintained at 4 °C during sorting.

### scRNA-seq library preparation and sequencing

Preparation of scRNA-seq libraries was carried out using a Chromium Single Cell 3ʹ GEM, Library & Gel Bead Kit v3.1 (10x Genomics, #PN-1000075). We input between 10 and 20k cells per lane on a Chromium Chip G and followed the manufacturer’s protocol for transcriptomic library preparation. Transcriptomic libraries were sequenced on Illumina Novaseq X. We used paired-end sequencing on Illumina platforms, targeting 50,000 reads per cell for cDNA libraries.

### STICR barcode library preparation

STICR barcodes were subamplified from each 10X cDNA library using the Q5 Hot Start High Fidelity 2× master mix (NEB, M0494). In brief, 10 μl of cDNA was used as template in a 50 μl PCR reaction containing STICR barcode read 1 and 2 primers (0.5 μM, each) targeting the region immediately upstream of the STICR viral index/barcode as well as the partial Illumina Read1 sequence added during cDNA library preparation (Table S5), using the following program: 1, 98 °C, 30 s; 2, 98 °C, 10 s; 3, 62 °C, 20 s; 4, 72 °C, 10 s; 5, repeat steps 2–4 11 times; 6, 72 °C, 2 min; 7, 4 °C, hold. Following PCR amplification, a 0.8–0.6 dual-sided size selection was performed using Ampure XP beads (Beckman Coulter, #A63881). Barcode libraries were sequenced on the NextSeq 2000. We used paired-end sequencing on Illumina platforms, targeting 5,000 reads per cell for barcode libraries.

### scRNA-seq analysis for mouse data

10X transcriptomic libraries were processed using CellRanger (version 7.2.0) and were aligned to the mouse transcriptome mm10-2020-A. Cellranger outputs were processed using Seurat v4.3.0.9002. Datasets were processed to remove cells with fewer than 1,000 or greater than 100,000 reads, fewer than 500 or greater than 10,000 genes, or more than 20% of reads corresponding to mitochondrial genes. Doublets were identified and removed by using DoubletFinder (version 2.0.6). Libraries were integrated using Seurat’s SCTransform and Harmony v0.1.162. To identify clusters, we constructed a shared-nearest neighbor graph based on Harmony embeddings via the “FindNeighbors” function to use as input to the SLM algorithm, implemented through the “FindClusters” function in Seurat. Differential expression analysis was conducted using Seurat’s “FindMarkers” and “FindAllMarkers” function, and gene expression in clusters was manually analyzed for known markers of given cell types. Iterative clustering to identify subtypes of cells was performed by subsetting the full Seurat object to only cells that were present in desired clusters, then re-running Harmony integration, cluster identification, and dimensional reduction. Cell identities determined through analysis of iterative clusters were reassigned to cells in the full Seurat object.

### STICR barcode analysis

STICR barcode analysis was performed using custom scripts, as previously described in Delgado et al30. First, BBMap (BBMap, Bushnell B; sourceforge.net/projects/bbmap/) was used to remove low-quality reads and then extract reads containing STICR barcode sequences. Then, BBMap was used to extract individual STICR barcode fragments that were then aligned to our predefined fragment reference sets using Bowtie47 (version 5.2.1), allowing for up to two mismatches per fragment. Aligned STICR barcodes were compiled into a file containing their corresponding 10x cell barcode and 10x UMI sequences using Awk. Finally, UMI-tools48 (version 0.5.1) was used to remove duplicate STICR barcode/cell barcode (CBC) reads by UMI, allowing for 1-bp mismatches in the UMI. STICR barcodes/CBC) pairings with at least five distinct UMIs were retained. Cells with a single STICR barcode meeting this criterion were retained for clonal analysis. Possible instances of STICR barcode superinfection (multiple STICR barcodes per starting progenitor) were identified by calculating jaccard similarity indexes of all STICR barcodes pairings found to co-occur within a single cell. Those pairings with a jaccard similarity index of 0.55 or greater that occurred in 10 cells or more were considered to be a valid superinfection clone and retained for clonal analysis. Cells that contained multiple STICR barcodes that were not determined to be valid superinfections were further analysed for the relative abundance of individual STICR barcodes. Cells that contained a ‘dominant’ STICR barcode with five or more times the number of barcode counts compared with the next most abundant STICR barcode, as determined by UMI, were retained and assigned that dominant barcode. Those cells that did not contain barcodes meeting these criteria were not considered for clonal analysis. To ensure high fidelity of barcode assignments to cells, several steps were taken with custom downstream processing steps. CBC/barcode combinations with fewer than 10 reads were removed from downstream analysis. The final filtering step was to remove barcode calls with fewer than 3 UMIs. Barcodes and barcode UMI counts were mapped to cells in the Seurat object by aligning on the cell barcode.

### DNA extraction for genotype array data

DNA Extraction for SNP genotyping assay was performed using PureLink Genomic DNA extraction kit (CAT; K182001) extraction using protocols from the manufacturer. Sample quality and concentrations were measured on a NanoDrop One spectrophotometer. All samples that passed QC were > 1.6-1.9 for 260/280 and contained a total of > 2000 ng of DNA. Isolated DNA from samples was sequenced using the iSCAN instrument (Illumina) for the Global Screening Array (GSAv2) chip at the QB3 facility at the University of California, Berkeley.

### snMultiome library preparation and sequencing

Dissected Putamen samples were obtained from donors within the Autism Brain Net (N=86),). Details such as postmortem interval, sex, age, and diagnosis were used to define a cohort with largely comparable properties. We were not able to confirm the genotype and SNP for donor #50 and removed the donor from further analysis. One more donor was filtered in the quality control step of single cell multiome data.

#### Nuclei Isolation and sn-multiome generation

Experimenters isolating nuclei from all donors were blinded to sample genotype and sex. All donor tissue samples were shipped with unique shipping IDs from Autism Brain Net, stored until nuclei or DNA extraction for SNP genotyping. Nuclei from the frozen tissue were extracted using an adapted protocol from the Kriegstein lab at UCSF. Frozen tissue blocks from individuals were dissected and cut into groups of ten to twelve on dry ice. An equal amount of each sample was obtained using this method and added to achieve a final weight of 50-100mg of pooled tissue in a shared tube of frozen tissue. The resulting tube contained a pool of frozen tissue with equal weight representation from each sample. The tube containing the pool of frozen tissue was then homogenized using a pre-chilled Dounce and a homogenization buffer consisting of .2653 M Sucrose, 26.6 mM KCl, 5.31 mM MgCl2, 21.2 mM Tricine-KOH pH 7.8, Ultrapure H2O, 1 mM DTT, .5 mM Spermidine, .15 mM Spermine, .3% NP40, 1X Protease Inhibitor, and .6 U/uL RNAse Inhibitor. The resulting homogenized mixture was then centrifuged to pellet the sample, and the supernatant was removed. The nuclei within the sample were isolated using an iodixanol gradient, through which varying iodixanol concentrations separated the nuclei band from the debris and myelin. Post-centrifugation, the nuclear band was extracted from the gradient and diluted with wash buffer containing 10 mM Tris-HCl pH 7.4, 10 mM NaCl, 3 mM MgCl2, 1% BSA, 0.1% Tween-20, 1 mM DTT, 0.6 U/uL RNAse Inhibitor, and ultrapure H2O. The resulting diluted sample was then counted manually using a hemocytometer and Trypan Blue staining. The solution was then centrifuged, and the nuclear pellet was resuspended to a concentration of 10,000-12,000 nuclei/uL. The final nuclei were suspended in the recommended Diluted Nuclei Buffer containing 1X Nuclei Buffer (PN-2000207, 10x Genomics), 1 mM DTT, 1 U/uL RNase Inhibitor, and Nuclease-Free H2O as recommended. The suspended nuclei resulting from the nuclei isolation protocol were split into four 10x Genomics Multiome wells with around 40,000 nuclei input per well. The target nuclei recovery was about 18,000-20,000 nuclei per well.

The following steps were followed per Multiome protocol using 10x Genomics’ Chromium Next GEM Single Cell Multiome ATAC + Gene Expression User Guide Rev E. The resulting Gene Expression and ATAC libraries underwent quality control and quantification using a High Sensitivity DNA Bioanalyzer 2100 (Agilent) to ensure adequate concentration and curve quality before sequencing. Equal moral ratios of all libraries were sequenced through the UCSF Center for Advanced Technology (CAT) and the Fabrication and Design Center. The Gene Expression libraries were sequenced on a Novaseq X 10B/25B flow cell under the following parameters: Read 1 of 28 bp, Index 1 of 10 bp, Index 2 of 10 bp, and Read 2 of 90 bp. The targeted reads per cell for the Gene Expression libraries were 25,000 reads. The ATAC libraries were sequenced on a Novaseq X 25B flow cell under the following parameters: Read 1 of 151 bp, Index 1 of 12 bp, Index 2 of 24 bp, and Read 2 of 151 bp. ATAC libraries were targeted for 30,000 reads. All sequenced libraries utilized the recommended 1% PhiX spike-in. All libraries were sequenced one or more times to achieve a final satisfactory read depth.

### Preprocessing of scMultiome data

#### Read preprocessing, alignment, and count generation

Read preprocessing, alignment, and count generation were done using 10x CellRanger-ARC version 2.0.2 with genome reference 2020-A (July 7, 2020), which uses GRCh38 primary assembly from Ensembl 98 and gene annotations/structure from GENCODE v32 primary assembly. CellRanger-ARC was run per 10x well to identify well-specific quality control metrics.

#### Genotype array QC

Genotype array data (GSAv2, GRCh38) were QCed and imputed to the TOPMED imputation panel^105,106^. Genotype QC was conducted using PLINK v1.90b7 (16 Jan 2023). Samples were checked for discordant sex (1 sample was ambiguous), high heterozygosity (± 3 SD from the mean), high missingness (± 3 SD from the mean), and relatedness. One sample failed all QC measures, and an additional three were found to have high heterozygosity; however, this was explained by having “admixed” ancestry or ancestry divergent from the majority of samples through principal component analysis (PCA) done with 1,000 Genomes^107^. Two samples were identified as identical. There was no indication in the phenotype data that these two samples were twins (different year of birth), indicating a sample mix-up. These were removed. SNPs were filtered for high call rate (≥ 0.98), and Hardy-Weinberg Equilibrium (p < 1×10-6). Ambiguous (aka palindromic A/T, G/C) SNPs and monomorphic SNPs (i.e., SNPs where everyone is homozygous), were removed. SNPs were prepared for imputation against TOPMED using a check tool by Will Rayner and can be found here: https://www.chg.ox.ac.uk/∼wrayner/tools/. The script compares strand, alleles, position, reference and alternate (REF/ALT) assignments and frequency differences between the data and the reference panel. It subsequently updates strand, position and REF/ALT assignment to match the reference and removed ambiguous SNPs if MAF > 0.4, SNPs with differing alleles, SNPs with > 0.2 allele frequency difference and SNPs not in the reference panel. Post imputation, SNPs were filtered for MAF ≥ 5% and imputation R^2^ ≥ 0.95.

#### PCA

Principal components were generated through PCA using PLINK v2.00a4.2LM (31 May 2023). QC-ed SNPs were subset to those overlapping with unrelated individuals in 1,000 Genomes (up to 2nd degree relatives were removed). We used linkage disequilibrium (LD)-pruned SNPs (pairwise r^2^ <0.2 in batches of 50 SNPs with sliding windows of 5) with MAF > 5% and removed 24 regions with high or long-range LD, including the HLA, leaving 294,312 variants for PCA.

#### Donor assignment

Donor assignment of barcodes was done using demuxlet^108^ with modified code from Vincent Gardieux (https://github.com/VincentGardeux/demuxlet.git) to avoid issues with random access memory (RAM) when using a large number of SNPs. As input to demuxlet, we included QC-ed and imputed genotype data (See “Genotype QC”) and bam files output from 10x CellRanger-ARC version 2.0.2 (See “Read preprocessing, alignment and count generation”).

#### Ambient RNA

We used CellBender^109^ v0.3.0 to identify barcodes likely tagging ambient RNA rather than nuclei.

### Quality control of scMultiome data

Nuclei in each multiome well were filtered after removing ambient RNA as determined by CellBender, and singlets as called by donor assignment (demuxlet), according to the following criteria: Percent mitochondrial reads < 8%, Blacklist ratio < 5%, 2% < transcription start-site (TSS) enrichment < 15%, ln(gene counts + 1) within 4 median-absolute deviations (MADs), ln(RNA counts + 1) within 4 MADs, ln(ATAC counts + 1) within 4 MADs, Promoter ratio within 4 MADs, Percent reads in peaks within 5 MADs, Nucleosome signal within 5 MADs.

### Cell type annotation

We normalised the RNA gene count matrices using SCTransform in Seurat^110^ v5. We then used Harmony^111^ (v0.1.1) to correct batch effects across wells and shared donors. Integration was performed on the top 30 principal components from the SCT assay using the default parameters: theta = c(2,2), lambda = c(1,1), and sigma = 0.1. Harmony embeddings were used for downstream UMAP visualization and Louvain clustering (resolution = 1) based on a shared nearest neighbor graph constructed from top 30 dimensions. Broad cell types were annotated manually based on canonical marker gene expression. Subtypes of Medium Spiny Neurons were identified by reference mapping to the Cell Type Taxonomies (dorsal striatum) of the human basal ganglia using MapMyCells (RRID: SCR_024672).

### Cell proportion analysis

Cell-type composition was performed at the donor level using the Propeller R package (v1.12.0). Donors with fewer than 300 cells or containing less than half of the annotated cell types (≥5 cells per cluster) were excluded. Unannotated clusters (“UN”) were omitted. Cell-type proportions were computed using the getTransformedProps() function from Propeller, which calculates the relative abundance of each cell type per donor and applies a logit transformation to stabilize variance across compositions. Differences in cell-type proportions between case and control donors were assessed using the Wilcoxon rank-sum test, and P values were corrected for multiple testing using the Benjamini–Hochberg method.

### Differential gene expression analysis

We used *dreamlet* v1.0.3 for differential gene expression after pseudobulking cells by donor per cell type. We compared donors with (N = 14) and without (N = 33) a genetic diagnosis to controls (N = 37), controlling for RNA Integrity Number, Post-Mortem Interval, sex, age, median percent mitochondrial reads, number of genes expressed, tissue group and top two genetic principal components. Covariates were chosen on the basis of variance explained (> 1 %) or correlation with case status (p value 0.05 with 17 correlations tested), ensuring no two covariates were correlated with r^2^ ≥ 0.6 to avoid collinearity. Differential expression analysis between donors without a diagnosis and controls was done among male donors only (with sex covariate removed), as only one donor without a genetic diagnosis was female. Genes were filtered by default *dreamlet* settings (min.cells = 5, min.count = 5, min.samples = 4, min.prop = 0.4). To check for inflation of results, we ran differential expression analysis with shuffled case status labels, and plotted quantile-quantile (QQ) plots of the resultant p values per cell type (Figure S5). As only one genetically undiagnosed donor was female, we limited our analysis to male donors and omitted sex from the model for DGE analysis of donors without a genetic diagnosis.

### Biological process enrichment analysis

Functional enrichment analysis was performed using the “clusterProfiler” R package^112,113^ (v4.10.1) with default parameters. The functional annotations were obtained from Kyoto Encyclopedia of Genes and Genomes (KEGG) pathway^114^ and terms of biological processes in Gene Ontology (GO)^115^.

### Peak calling

Peak calling was performed following the pycisTopic (version 2.0a0) workflow. Cell-type labels from the annotation using scRNA-seq data were used to generate pseudobulk accessibility profiles for each cell type and brain region. Peaks were called using MACS2 (version 2.2.9.1) with the parameters --shift 73 --extsize 146 --qvalue 0.05, yielding high-confidence open chromatin regions per cell type. To generate a unified set of accessible regions, we applied the TGCA iterative peak filtering approach implemented in pycisTopic to derive species-specific consensus peaks. Briefly, each peak was extended by 250 bp in both directions from its summit, and less significant peaks that overlapped with more significant ones were iteratively removed. This process resulted in a final set of 1,166,673 consensus peaks, obtained using the “get_consensus_peaks” function in pycisTopic after removing ENCODE blacklisted regions.

### Peak to gene linkage

#### SEACells analysis

To enhance the statistical power of peak-to-gene inference, we first generated metacells using the SEACells algorithm^116^. SEACells defines groups of transcriptionally similar cells to enhance the signal sensitivity of both RNA and ATAC modalities. Following batch-effect correction, the harmony embedding space was used to construct the initial kernel matrix. To improve computational efficiency, SEACells was applied independently to each cell type within the batch-corrected harmony space. For each cell type, approximately 100 single cells were aggregated to form one metacell, while smaller clusters contained a minimum of 20 metacells. The SEACells function was executed with parameters n_waypoint_eigs = 10 and convergence_epsilon = 1e-5, and the model was fitted for a maximum of 50 iterations. This procedure resulted in the identification of 3,369 metacells across the dataset. Subsequently, average normalized RNA and ATAC profiles were computed for each metacell using the “AverageExpression” function in Seurat, providing inputs for downstream peak–gene association analyses.

#### Peak-to-gene linkage

Peak-to-gene associations were inferred using the LinkPeaks function implemented in the Signac package^117^ (version 1.14.0). Normalized matrices stored in the data slot were used for both the peak and RNA assays to ensure consistency across modalities. For each gene, LinkPeaks computes the correlation coefficient between gene expression and chromatin accessibility of peaks located within a defined genomic window around the TSS. The function further estimates an expected correlation coefficient for each peak based on its GC content, accessibility, and length, and then calculates a z-score and P-value for each observed correlation. Peaks with significant positive correlations (P < 0.05) and correlation scores greater than 0.4 were retained, while other parameters were set to their default values.

### Motif annotation and enrichment analysis

Motif annotation and enrichment were performed using Signac (v1.14) with position frequency matrices from JASPAR2020. Motifs were added to the ATAC assay using the AddMotifs() function and the hg38 reference genome (BSgenome.Hsapiens.UCSC.hg38). For the motif enrichment analysis, peaks linked to differentially expressed genes (FDR < 0.05, |log₂FC| > 0.2) were defined as the foreground, and peaks linked to all genes expressed in corresponding cell types served as the background. Enrichment was assessed with FindMotifs(), which applies a hypergeometric test to identify transcription factor motifs overrepresented in the foreground relative to background peaks. P values were adjusted using the Benjamini–Hochberg method.

### Gene Regulatory Network Analysis

We applied the SCENIC+^54^ workflow (v1.0a1) to construct gene regulatory networks (GRNs) from single-nucleus multiomic data. Because the full dataset contained over 300,000 nuclei, which is computationally demanding for GRN reconstruction, we subsampled one-tenth of all nuclei while retaining at least 100 cells from each sub-cell type to preserve transcriptional diversity. We next performed topic modeling using pycisTopic, which decomposes the accessibility matrix into latent “topics”, sets of co-accessible regions that may correspond to underlying regulatory programs. The optimal topic number (45) was determined by maximizing the model log-likelihood. To define candidate enhancer regions, three complementary strategies were used together: (1) binarization of topic weights using the Otsu method; (2) selection of the top 3,000 regions per topic; an (3) identification of differentially accessible peaks (log2[FC] > 0.5, adjusted P < 0.05) using the Wilcoxon rank-sum test on imputed accessibility values.

These candidate enhancers were subjected to motif enrichment analysis using pycistarget and a discrete element method (DEM)-based approach to identify overrepresented TF-binding motifs, thereby assigning putative TF–region relationships. A custom cisTarget database was built for motif annotation on the 1,166,673 consensus peaks by using “create_cistarget_motif_databases.py” with default parameters: --bgpadding 1000. TF–region– gene regulation network was inferred by using the SCENIC+ snakemake pipeline with default parameters. eRegulons were then defined as TF–region–gene triplets, comprising a TF, all regions enriched for its motif, and the genes linked to those regions based on co-accessibility and correlation patterns. For each eRegulon, specificity scores were computed using the Regulon Specificity Score (RSS) algorithm, based on Area Under the Curve (AUC) enrichment values for either region- or gene-based eRegulons. The top eRegulons per cell type were visualized as cell-type–specific regulatory modules.

### Estimating GWAS enrichment using Multiome peaks

To estimate the enrichment of various complex traits, we used LDSC^118^ (v2.0.1). GWAS summary statistics for the following traits were processed: autism spectrum disorder, age of walking, schizophrenia, Alzheimer’s disease, anorexia nervosa, educational attainment, epilepsy, intelligence, major depressive disorder, multiple sclerosis, obsessive-compulsive disorder, Parkinson’s disease, Tourette’s syndrome, problematic alcohol use, bipolar disorder, panic disorder, attention deficit hyperactivity disorder, developmental disorders, generalized epilepsy, childhood epilepsy, height, substance use disorder, post-traumatic stress disorder, frontotemporal lobar degeneration, and amyotrophic lateral sclerosis. We first harmonized the GWAS summary statistics using MungeSumstats v1.14.1, which included removing indels and excluding SNPs with imputation information scores below 0.7 when available. Next, we performed gene-level association analyses using MAGMA v1.08, as implemented in FUMA v1.5.2, with a ±1 kb window upstream and downstream of each gene to assign SNPs. MAGMA tested each protein-coding gene containing assigned SNPs, and genes were then ranked by their MAGMA p-values to identify the top trait-associated genes.

Cell-type-specific peaks were also lifted over to hg19 using the UCSD liftover tool (chain file: https://hgdownload.soe.ucsc.edu/goldenPath/hg38/liftOver/hg38ToHg19.over.chain.gz) and formatted for LDSC analysis with make_annotation.py. Linkage disequilibrium (LD) scores were then computed for each annotation set using ldsc.py. To account for multiple testing, false discovery rate (FDR) correction was applied to the enrichment P-values.

### GWAS fine-mapping

For GWAS locus selection, we first identified significant risk loci from ASD and ADHD GWAS datasets. We then selected SNPs located within gene-correlated CREs (peak-to-gene correlation > 0.3 and FDR < 0.05) as lead positions. We selected a neighborhood of around 1.5Mb that centers on the lead position. After fine-mapping these loci using ASD and ADHD summary statistics, we identified two loci that overlapped with cell-type–specific MSN D1 Striosome CREs. For genetic fine-mapping, we incorporated peak enrichment scores as prior weights and applied the Sum of Single-Effects (SuSiE, v0.14.2) regression model to infer risk variants. By integrating epigenetic information from our snATAC peaks, these priors enhanced the accuracy of causal variant detection.

For SNPs within ATAC peaks, we set the prior weights to 5 for SuSiE to prioritize SNPs with strong ATAC signals. For SNPs outside ATAC peaks, we set the prior weights to 0.1. CREs were defined by peak-to-gene correlations > 0.4 and FDR < 0.05, and these were used to guide SuSiE fine-mapping. Fine-mapping was performed using the susie_rss function in SuSiE, using ASD GWAS summary statistics. SuSiE computes the posterior inclusion probability (PIP) for each SNP, representing the probability that a variant is causally associated with the trait.

Linkage disequilibrium (LD) information was derived from 503 European samples from the 1000 Genomes Project (1KG). The 1KG Phase 3 dataset, obtained from the PLINK 2.0 Resources page (https://www.cog-genomics.org/plink/2.0/resources) was used as the reference panel. We selected European samples based on the 1KG super-population metadata and excluded duplicate or ambiguous SNPs. Quality control was performed in PLINK, retaining only SNPs with a genotyping rate ≥99% and minor allele frequency ≥0.005. Samples with >5% missing genotypes and markers failing the Hardy–Weinberg equilibrium test > 1e-6 were excluded. The final reference panel comprised 633 samples (328 females, 305 males) European samples sequenced genotyped at 8,902,823 variants. The coordinates of risk SNPs from ADHD and SCZ were lifted over to hg38 using the UCSD liftover tool (chain file: https://hgdownload.soe.ucsc.edu/goldenPath/hg19/liftOver/hg19ToHg38.over.chain.gz)

For visualization, we integrated multiple layers of information, including ATAC signal tracks, CRE–gene links, SuSiE PIP values, and gene tracks across predefined genomic regions. We plotted all variants, highlighted variants with PIP > 0.5, and differentiated variant markers by shape when located within a CRE. CRE–gene loops were color-coded according to their correlation coefficient values.

## Statistics and Reproducibility

The number of brains/samples/donors collected from each time point are listed in the supplementary table. No prior statistics were used to determine the number of donors.

## Data availability

scRNA-seq transcriptomic data and STICR barcode data are available at the Gene Expression Omnibus (GEO; https://www.ncbi.nlm.nih.gov/geo/) under accession number GSExxxx. An interactive browser of single-cell data will be found at the University of California, Santa Cruz (UCSC) cell browser.

## Code availability

Custom codes used in this study are available at the following GitHub repository: https://github.com/NOW-Lab/STICR

## REFERENCE

1. Hodges, H., Fealko, C., and Soares, N. (2020). Autism spectrum disorder: definition, epidemiology, causes, and clinical evaluation. Transl. Pediatr. 9, S55–S65.

2. Hirota, T., and King, B.H. (2023). Autism spectrum disorder: A review. JAMA 329, 157–168.

3. Colvert, E., Tick, B., McEwen, F., Stewart, C., Curran, S.R., Woodhouse, E., Gillan, N., Hallett, V., Lietz, S., Garnett, T., et al. (2015). Heritability of autism spectrum disorder in a UK population-based twin sample. JAMA Psychiatry 72, 415–423.

4. Sandin, S., Lichtenstein, P., Kuja-Halkola, R., Hultman, C., Larsson, H., and Reichenberg, A. (2017). The heritability of autism spectrum disorder. JAMA 318, 1182–1184.

5. Iossifov, I., O’Roak, B.J., Sanders, S.J., Ronemus, M., Krumm, N., Levy, D., Stessman, H.A., Witherspoon, K.T., Vives, L., Patterson, K.E., et al. (2014). The contribution of de novo coding mutations to autism spectrum disorder. Nature 515, 216–221.

6. Satterstrom, F.K., Kosmicki, J.A., Wang, J., Breen, M.S., De Rubeis, S., An, J.-Y., Peng, M., Collins, R., Grove, J., Klei, L., et al. (2020). Large-scale exome sequencing study implicates both developmental and functional changes in the neurobiology of autism. Cell 180, 568–584.e23.

7. Fu, J.M., Satterstrom, F.K., Peng, M., Brand, H., Collins, R.L., Dong, S., Wamsley, B., Klei, L., Wang, L., Hao, S.P., et al. (2022). Rare coding variation provides insight into the genetic architecture and phenotypic context of autism. Nat. Genet. 54, 1320–1331.

8. Weiss, L.A., Shen, Y., Korn, J.M., Arking, D.E., Miller, D.T., Fossdal, R., Saemundsen, E., Stefansson, H., Ferreira, M.A.R., Green, T., et al. (2008). Association between microdeletion and microduplication at 16p11.2 and autism. N. Engl. J. Med. 358, 667–675.

9. Niarchou, M., Chawner, S.J.R.A., Doherty, J.L., Maillard, A.M., Jacquemont, S., Chung, W.K., Green-Snyder, L., Bernier, R.A., Goin-Kochel, R.P., Hanson, E., et al. (2019). Psychiatric disorders in children with 16p11.2 deletion and duplication. Transl. Psychiatry 9, 8.

10. Voineagu, I., Wang, X., Johnston, P., Lowe, J.K., Tian, Y., Horvath, S., Mill, J., Cantor, R.M., Blencowe, B.J., and Geschwind, D.H. (2011). Transcriptomic analysis of autistic brain reveals convergent molecular pathology. Nature 474, 380–384.

11. Jia, Y., Zhao, C., Wang, Q., Shu, C., Feng, X., Song, F., and Zhang, J. (2014). A genetically modified broad-spectrum strain of Bacillus thuringiensis toxic against Holotrichia parallela, Anomala corpulenta and Holotrichia oblita. World J. Microbiol. Biotechnol. 30, 595–603.

12. Forrest, M.P., Parnell, E., and Penzes, P. (2018). Dendritic structural plasticity and neuropsychiatric disease. Nat. Rev. Neurosci. 19, 215–234.

13. Xu, X., Wells, A.B., O’Brien, D.R., Nehorai, A., and Dougherty, J.D. (2014). Cell type-specific expression analysis to identify putative cellular mechanisms for neurogenetic disorders. J. Neurosci. 34, 1420–1431.

14. Kelvington, B.A., Kim, J., Fair, R., Gaine, M.E., and Abel, T. (2025). Complement contributes to hyperactive behavior in the 16p11.2 hemideletion mouse model. bioRxivorg. 10.1101/2025.08.21.671537.

15. Evans, M.M., Kim, J., Abel, T., Nickl-Jockschat, T., and Stevens, H.E. (2024). Developmental disruptions of the dorsal striatum in autism spectrum disorder. Biol. Psychiatry 95, 102–111.

16. Peça, J., Feliciano, C., Ting, J.T., Wang, W., Wells, M.F., Venkatraman, T.N., Lascola, C.D., Fu, Z., and Feng, G. (2011). Shank3 mutant mice display autistic-like behaviours and striatal dysfunction. Nature 472, 437–442.

17. Fuccillo, M.V. (2016). Striatal circuits as a common node for autism pathophysiology. Front. Neurosci. 10, 27.

18. Araujo, D.J., Anderson, A.G., Berto, S., Runnels, W., Harper, M., Ammanuel, S., Rieger, M.A., Huang, H.-C., Rajkovich, K., Loerwald, K.W., et al. (2015). FoxP1 orchestration of ASD-relevant signaling pathways in the striatum. Genes Dev. 29, 2081–2096.

19. Langen, M., Bos, D., Noordermeer, S.D.S., Nederveen, H., van Engeland, H., and Durston, S. (2014). Changes in the development of striatum are involved in repetitive behavior in autism. Biol. Psychiatry 76, 405–411.

20. Chang, J., Gilman, S.R., Chiang, A.H., Sanders, S.J., and Vitkup, D. (2015). Genotype to phenotype relationships in autism spectrum disorders. Nat. Neurosci. 18, 191–198.

21. Cataldi, S., Stanley, A.T., Miniaci, M.C., and Sulzer, D. (2022). Interpreting the role of the striatum during multiple phases of motor learning. FEBS J. 289, 2263–2281.

22. Cox, J., and Witten, I.B. (2019). Striatal circuits for reward learning and decision-making. Nat. Rev. Neurosci. 20, 482–494.

23. Sato, W., Kubota, Y., Kochiyama, T., Uono, S., Yoshimura, S., Sawada, R., Sakihama, M., and Toichi, M. (2014). Increased putamen volume in adults with autism spectrum disorder. Front. Hum. Neurosci. 8, 957.

24. Estes, A., Shaw, D.W.W., Sparks, B.F., Friedman, S., Giedd, J.N., Dawson, G., Bryan, M., and Dager, S.R. (2011). Basal ganglia morphometry and repetitive behavior in young children with autism spectrum disorder. Autism Res. 4, 212–220.

25. Abbott, A.E., Linke, A.C., Nair, A., Jahedi, A., Alba, L.A., Keown, C.L., Fishman, I., and Müller, R.-A. (2018). Repetitive behaviors in autism are linked to imbalance of corticostriatal connectivity: a functional connectivity MRI study. Soc. Cogn. Affect. Neurosci. 13, 32–42.

26. Sanders, S.J., Murtha, M.T., Gupta, A.R., Murdoch, J.D., Raubeson, M.J., Willsey, A.J., Ercan-Sencicek, A.G., DiLullo, N.M., Parikshak, N.N., Stein, J.L., et al. (2012). De novo mutations revealed by whole-exome sequencing are strongly associated with autism. Nature 485, 237–241.

27. Kim, J., Vanrobaeys, Y., Kelvington, B., Peterson, Z., Baldwin, E., Gaine, M.E., Nickl-Jockschat, T., and Abel, T. (2024). Dissecting 16p11.2 hemi-deletion to study sex-specific striatal phenotypes of neurodevelopmental disorders. Mol Psychiatry 29, 1310–1321.

28. Shin, D., Kim, C.N., Ross, J., Hennick, K.M., Wu, S.-R., Paranjape, N., Leonard, R., Wang, J.C., Keefe, M.G., Pavlovic, B.J., et al. (2024). Thalamocortical organoids enable in vitro modeling of 22q11.2 microdeletion associated with neuropsychiatric disorders. Cell Stem Cell 31, 421–432.e8.

29. Perez, Y., Velmeshev, D., Wang, L., White, M., Siebert, C., Baltazar, J., Dutton, N.G., Wang, S., Haeussler, M., Chamberlain, S., et al. (2023). Single cell analysis of dup15q syndrome reveals developmental and postnatal molecular changes in autism. bioRxiv. 10.1101/2023.09.22.559056.

30. Blumenthal, I., Ragavendran, A., Erdin, S., Klei, L., Sugathan, A., Guide, J.R., Manavalan, P., Zhou, J.Q., Wheeler, V.C., Levin, J.Z., et al. (2014). Transcriptional consequences of 16p11.2 deletion and duplication in mouse cortex and multiplex autism families. Am J Hum Genet 94, 870–883.

31. Delgado, R.N., Allen, D.E., Keefe, M.G., Mancia Leon, W.R., Ziffra, R.S., Crouch, E.E., Alvarez-Buylla, A., and Nowakowski, T.J. (2022). Individual human cortical progenitors can produce excitatory and inhibitory neurons. Nature 601, 397–403.

32. Zhang, M., Pan, X., Jung, W., Halpern, A.R., Eichhorn, S.W., Lei, Z., Cohen, L., Smith, K.A., Tasic, B., Yao, Z., et al. (2023). Molecularly defined and spatially resolved cell atlas of the whole mouse brain. Nature 624, 343–354.

33. Yao, Z., van Velthoven, C.T.J., Kunst, M., Zhang, M., McMillen, D., Lee, C., Jung, W., Goldy, J., Abdelhak, A., Aitken, M., et al. (2023). A high-resolution transcriptomic and spatial atlas of cell types in the whole mouse brain. Nature 624, 317–332.

34. Tinterri, A., Menardy, F., Diana, M.A., Lokmane, L., Keita, M., Coulpier, F., Lemoine, S., Mailhes, C., Mathieu, B., Merchan-Sala, P., et al. (2018). Active intermixing of indirect and direct neurons builds the striatal mosaic. Nat. Commun. 9, 4725.

35. Kelly, S.M., Raudales, R., He, M., Lee, J.H., Kim, Y., Gibb, L.G., Wu, P., Matho, K., Osten, P., Graybiel, A.M., et al. (2018). Radial glial lineage progression and differential intermediate progenitor amplification underlie striatal compartments and circuit organization. Neuron 99, 345–361.e4.

36. Bandler, R.C., Vitali, I., Delgado, R.N., Ho, M.C., Dvoretskova, E., Ibarra Molinas, J.S., Frazel, P.W., Mohammadkhani, M., Machold, R., Maedler, S., et al. (2022). Single-cell delineation of lineage and genetic identity in the mouse brain. Nature 601, 404–409.

37. Phan, B.N., Ray, M.H., Xue, X., Fu, C., Fenster, R.J., Kohut, S.J., Bergman, J., Haber, S.N., McCullough, K.M., Fish, M.K., et al. (2024). Single nuclei transcriptomics in human and non-human primate striatum in opioid use disorder. Nat. Commun. 15, 878.

38. Tran, M.N., Maynard, K.R., Spangler, A., Huuki, L.A., Montgomery, K.D., Sadashivaiah, V., Tippani, M., Barry, B.K., Hancock, D.B., Hicks, S.C., et al. (2021). Single-nucleus transcriptome analysis reveals cell-type-specific molecular signatures across reward circuitry in the human brain. Neuron 109, 3088–3103.e5.

39. Garma, L.D., Harder, L., Barba-Reyes, J.M., Marco Salas, S., Díez-Salguero, M., Nilsson, M., Serrano-Pozo, A., Hyman, B.T., and Muñoz-Manchado, A.B. (2024). Interneuron diversity in the human dorsal striatum. Nat. Commun. 15, 6164.

40. Escartin, C., Galea, E., Lakatos, A., O’Callaghan, J.P., Petzold, G.C., Serrano-Pozo, A., Steinhäuser, C., Volterra, A., Carmignoto, G., Agarwal, A., et al. (2021). Reactive astrocyte nomenclature, definitions, and future directions. Nat. Neurosci. 24, 312–325.

41. He, J., Kleyman, M., Chen, J., Alikaya, A., Rothenhoefer, K.M., Ozturk, B.E., Wirthlin, M., Bostan, A.C., Fish, K., Byrne, L.C., et al. (2021). Transcriptional and anatomical diversity of medium spiny neurons in the primate striatum. Curr. Biol. 31, 5473–5486.e6.

42. Phipson, B., Sim, C.B., Porrello, E.R., Hewitt, A.W., Powell, J., and Oshlack, A. (2022). Propeller: Testing for differences in cell type proportions in single cell data. Bioinformatics 38, 4720–4726.

43. Hoffman, G.E., Lee, D., Bendl, J., Prashant, N.M., Hong, A., Casey, C., Alvia, M., Shao, Z., Argyriou, S., Therrien, K., et al. (2024). Efficient differential expression analysis of large-scale single cell transcriptomics data using dreamlet. bioRxivorg. 10.1101/2023.03.17.533005.

44. Graybiel, A.M. (2008). Habits, rituals, and the evaluative brain. Annu. Rev. Neurosci. 31, 359–387.

45. Crittenden, J.R., and Graybiel, A.M. (2011). Basal Ganglia disorders associated with imbalances in the striatal striosome and matrix compartments. Front. Neuroanat. 5, 59.

46. Harris, J.J., Jolivet, R., and Attwell, D. (2012). Synaptic energy use and supply. Neuron 75, 762–777.

47. Rangaraju, V., Lauterbach, M., and Schuman, E.M. (2019). Spatially stable mitochondrial compartments fuel local translation during plasticity. Cell 176, 73–84.e15.

48. Stathopoulos, S., Gaujoux, R., Lindeque, Z., Mahony, C., Van Der Colff, R., Van Der Westhuizen, F., and O’Ryan, C. (2020). DNA methylation associated with mitochondrial dysfunction in a South African autism spectrum disorder cohort. Autism Res. 13, 1079– 1093.

49. Griffiths, K.K., and Levy, R.J. (2017). Evidence of mitochondrial dysfunction in autism: Biochemical links, genetic-based associations, and non-energy-related mechanisms. Oxid. Med. Cell. Longev. 2017, 4314025.

50. Galvez-Contreras, A.Y., Zarate-Lopez, D., Torres-Chavez, A.L., and Gonzalez-Perez, O. (2020). Role of oligodendrocytes and myelin in the pathophysiology of autism Spectrum Disorder. Brain Sci. 10, 951.

51. Closser, M., Guo, Y., Wang, P., Patel, T., Jang, S., Hammelman, J., De Nooij, J.C., Kopunova, R., Mazzoni, E.O., Ruan, Y., et al. (2022). An expansion of the non-coding genome and its regulatory potential underlies vertebrate neuronal diversity. Neuron 110, 70–85.e6.

52. Meuleman, W., Muratov, A., Rynes, E., Halow, J., Lee, K., Bates, D., Diegel, M., Dunn, D., Neri, F., Teodosiadis, A., et al. (2020). Index and biological spectrum of human DNase I hypersensitive sites. Nature 584, 244–251.

53. Fullard, J.F., Hauberg, M.E., Bendl, J., Egervari, G., Cirnaru, M.-D., Reach, S.M., Motl, J., Ehrlich, M.E., Hurd, Y.L., and Roussos, P. (2018). An atlas of chromatin accessibility in the adult human brain. Genome Res. 28, 1243–1252.

54. Bravo González-Blas, C., De Winter, S., Hulselmans, G., Hecker, N., Matetovici, I., Christiaens, V., Poovathingal, S., Wouters, J., Aibar, S., and Aerts, S. (2023). SCENIC+: single-cell multiomic inference of enhancers and gene regulatory networks. Nat. Methods 20, 1355–1367.

55. De Rubeis, S., He, X., Goldberg, A.P., Poultney, C.S., Samocha, K., Cicek, A.E., Kou, Y., Liu, L., Fromer, M., Walker, S., et al. (2014). Synaptic, transcriptional and chromatin genes disrupted in autism. Nature 515, 209–215.

56. Feliciano, P., Zhou, X., Astrovskaya, I., Turner, T.N., Wang, T., Brueggeman, L., Barnard, R., Hsieh, A., Snyder, L.G., Muzny, D.M., et al. (2019). Exome sequencing of 457 autism families recruited online provides evidence for autism risk genes. NPJ Genom. Med. 4, 19.

57. Alexander, E.J., Ghanbari Niaki, A., Zhang, T., Sarkar, J., Liu, Y., Nirujogi, R.S., Pandey, A., Myong, S., and Wang, J. (2018). Ubiquilin 2 modulates ALS/FTD-linked FUS-RNA complex dynamics and stress granule formation. Proc. Natl. Acad. Sci. U. S. A. 115, E11485–E11494.

58. Matthews, A.M., and Whiteley, A.M. (2025). UBQLN2 in neurodegenerative disease: mechanistic insights and emerging therapeutic potential. Biochem. Soc. Trans. 53, 823– 833.

59. Gui, A., Hollowell, A., Wigdor, E.M., Morgan, M.J., Hannigan, L.J., Corfield, E.C., Odintsova, V., Hottenga, J.-J., Wong, A., Pool, R., et al. (2024). Genome-wide association meta-analysis of age at onset of walking. medRxiv. 10.1101/2024.05.07.24306845.

60. Grove, J., Ripke, S., Als, T.D., Mattheisen, M., Walters, R.K., Won, H., Pallesen, J., Agerbo, E., Andreassen, O.A., Anney, R., et al. (2019). Identification of common genetic risk variants for autism spectrum disorder. Nat. Genet. 51, 431–444.

61. Trubetskoy, V., Pardiñas, A.F., Qi, T., Panagiotaropoulou, G., Awasthi, S., Bigdeli, T.B., Bryois, J., Chen, C.-Y., Dennison, C.A., Hall, L.S., et al. (2022). Mapping genomic loci implicates genes and synaptic biology in schizophrenia. Nature 604, 502–508.

62. Bellenguez, C., Küçükali, F., Jansen, I.E., Kleineidam, L., Moreno-Grau, S., Amin, N., Naj, A.C., Campos-Martin, R., Grenier-Boley, B., Andrade, V., et al. (2022). New insights into the genetic etiology of Alzheimer’s disease and related dementias. Nat. Genet. 54, 412– 436.

63. Watson, H.J., Yilmaz, Z., Thornton, L.M., Hübel, C., Coleman, J.R.I., Gaspar, H.A., Bryois, J., Hinney, A., Leppä, V.M., Mattheisen, M., et al. (2019). Genome-wide association study identifies eight risk loci and implicates metabo-psychiatric origins for anorexia nervosa. Nat. Genet. 51, 1207–1214.

64. Okbay, A., Wu, Y., Wang, N., Jayashankar, H., Bennett, M., Nehzati, S.M., Sidorenko, J., Kweon, H., Goldman, G., Gjorgjieva, T., et al. (2022). Polygenic prediction of educational attainment within and between families from genome-wide association analyses in 3 million individuals. Nat. Genet. 54, 437–449.

65. International League Against Epilepsy Consortium on Complex Epilepsies. Electronic address: (2014). Genetic determinants of common epilepsies: a meta-analysis of genome-wide association studies. Lancet Neurol. 13, 893–903.

66. Savage, J.E., Jansen, P.R., Stringer, S., Watanabe, K., Bryois, J., de Leeuw, C.A., Nagel, M., Awasthi, S., Barr, P.B., Coleman, J.R.I., et al. (2018). Genome-wide association meta-analysis in 269,867 individuals identifies new genetic and functional links to intelligence. Nat. Genet. 50, 912–919.

67. Major Depressive Disorder Working Group of the Psychiatric Genomics Consortium. Electronic address, and Major Depressive Disorder Working Group of the Psychiatric Genomics Consortium (2025). Trans-ancestry genome-wide study of depression identifies 697 associations implicating cell types and pharmacotherapies. Cell 188, 640–652.e9.

68. International Multiple Sclerosis Genetics Consortium (2019). Multiple sclerosis genomic map implicates peripheral immune cells and microglia in susceptibility. Science 365, eaav7188.

69. International Obsessive Compulsive Disorder Foundation Genetics Collaborative (IOCDF-GC) and OCD Collaborative Genetics Association Studies (OCGAS) (2018). Revealing the complex genetic architecture of obsessive-compulsive disorder using meta-analysis. Mol. Psychiatry 23, 1181–1188.

70. Kim, J.J., Vitale, D., Otani, D.V., Lian, M.M., Heilbron, K., 23andMe Research Team, Iwaki, H., Lake, J., Solsberg, C.W., Leonard, H., et al. (2024). Multi-ancestry genome-wide association meta-analysis of Parkinson’s disease. Nat. Genet. 56, 27–36.

71. Yu, D., Sul, J.H., Tsetsos, F., Nawaz, M.S., Huang, A.Y., Zelaya, I., Illmann, C., Osiecki, L., Darrow, S.M., Hirschtritt, M.E., et al. (2019). Interrogating the genetic determinants of Tourette’s syndrome and other tic disorders through genome-wide association studies. Am. J. Psychiatry 176, 217–227.

72. Zhou, H., Kember, R.L., Deak, J.D., Xu, H., Toikumo, S., Yuan, K., Lind, P.A., Farajzadeh, L., Wang, L., Hatoum, A.S., et al. (2023). Multi-ancestry study of the genetics of problematic alcohol use in over 1 million individuals. Nat. Med. 29, 3184–3192.

73. O’Connell, K.S., Koromina, M., van der Veen, T., Boltz, T., David, F.S., Yang, J.M.K., Lin, K.-H., Wang, X., Coleman, J.R.I., Mitchell, B.L., et al. (2025). Genomics yields biological and phenotypic insights into bipolar disorder. Nature 639, 968–975.

74. Demontis, D., Walters, G.B., Athanasiadis, G., Walters, R., Therrien, K., Nielsen, T.T., Farajzadeh, L., Voloudakis, G., Bendl, J., Zeng, B., et al. (2023). Genome-wide analyses of ADHD identify 27 risk loci, refine the genetic architecture and implicate several cognitive domains. Nat. Genet. 55, 198–208.

75. Huang, Q.Q., Wigdor, E.M., Malawsky, D.S., Campbell, P., Samocha, K.E., Chundru, V.K., Danecek, P., Lindsay, S., Marchant, T., Koko, M., et al. (2024). Examining the role of common variants in rare neurodevelopmental conditions. Nature 636, 404–411.

76. International League Against Epilepsy Consortium on Complex Epilepsies (2023). GWAS meta-analysis of over 29,000 people with epilepsy identifies 26 risk loci and subtype-specific genetic architecture. Nat. Genet. 55, 1471–1482.

77. Yengo, L., Vedantam, S., Marouli, E., Sidorenko, J., Bartell, E., Sakaue, S., Graff, M., Eliasen, A.U., Jiang, Y., Raghavan, S., et al. (2022). A saturated map of common genetic variants associated with human height. Nature 610, 704–712.

78. Hatoum, A.S., Colbert, S.M.C., Johnson, E.C., Huggett, S.B., Deak, J.D., Pathak, G., Jennings, M.V., Paul, S.E., Karcher, N.R., Hansen, I., et al. (2023). Multivariate genome-wide association meta-analysis of over 1 million subjects identifies loci underlying multiple substance use disorders. Nat. Ment. Health 1, 210–223.

79. Nievergelt, C.M., Maihofer, A.X., Atkinson, E.G., Chen, C.-Y., Choi, K.W., Coleman, J.R.I., Daskalakis, N.P., Duncan, L.E., Polimanti, R., Aaronson, C., et al. (2024). Genome-wide association analyses identify 95 risk loci and provide insights into the neurobiology of post-traumatic stress disorder. Nat. Genet. 56, 792–808.

80. Strom, N.I., Gerring, Z.F., Galimberti, M., Yu, D., Halvorsen, M.W., Abdellaoui, A., Rodriguez-Fontenla, C., Sealock, J.M., Bigdeli, T., Coleman, J.R., et al. (2025). Genome-wide analyses identify 30 loci associated with obsessive-compulsive disorder. Nat. Genet. 57, 1389–1401.

81. Pottier, C., Küçükali, F., Baker, M., Batzler, A., Jenkins, G.D., van Blitterswijk, M., Vicente, C.T., De Coster, W., Wynants, S., Van de Walle, P., et al. (2025). Deciphering distinct genetic risk factors for FTLD-TDP pathological subtypes via whole-genome sequencing. Nat. Commun. 16, 3914.

82. van Rheenen, W., van der Spek, R.A.A., Bakker, M.K., van Vugt, J.J.F.A., Hop, P.J., Zwamborn, R.A.J., de Klein, N., Westra, H.-J., Bakker, O.B., Deelen, P., et al. (2021). Common and rare variant association analyses in amyotrophic lateral sclerosis identify 15 risk loci with distinct genetic architectures and neuron-specific biology. Nat. Genet. 53, 1636–1648.

83. Kaplanis, J., Samocha, K.E., Wiel, L., Zhang, Z., Arvai, K.J., Eberhardt, R.Y., Gallone, G., Lelieveld, S.H., Martin, H.C., McRae, J.F., et al. (2020). Evidence for 28 genetic disorders discovered by combining healthcare and research data. Nature 586, 757–762.

84. Li, M., Santpere, G., Imamura Kawasawa, Y., Evgrafov, O.V., Gulden, F.O., Pochareddy, S., Sunkin, S.M., Li, Z., Shin, Y., Zhu, Y., et al. (2018). Integrative functional genomic analysis of human brain development and neuropsychiatric risks. Science 362, eaat7615.

85. Werling, D.M., Pochareddy, S., Choi, J., An, J.-Y., Sheppard, B., Peng, M., Li, Z., Dastmalchi, C., Santpere, G., Sousa, A.M.M., et al. (2020). Whole-genome and RNA sequencing reveal variation and transcriptomic coordination in the developing human prefrontal cortex. Cell Rep. 31, 107489.

86. Weissberg, O., and Elliott, E. (2021). The Mechanisms of CHD8 in Neurodevelopment and Autism Spectrum Disorders. Genes (Basel) 12. 10.3390/genes12081133.

87. O’Roak, B.J., Vives, L., Girirajan, S., Karakoc, E., Krumm, N., Coe, B.P., Levy, R., Ko, A., Lee, C., Smith, J.D., et al. (2012). Sporadic autism exomes reveal a highly interconnected protein network of de novo mutations. Nature 485, 246–250.

88. Portmann, T., Yang, M., Mao, R., Panagiotakos, G., Ellegood, J., Dolen, G., Bader, P.L., Grueter, B.A., Goold, C., Fisher, E., et al. (2014). Behavioral abnormalities and circuit defects in the basal ganglia of a mouse model of 16p11.2 deletion syndrome. Cell Rep. 7, 1077–1092.

89. Goto, S. (2023). Specificity of striatal dopamine D1 system in humans: implications for clinical use of D1 receptor-agonists in Parkinson’s disease. Front. Hum. Neurosci. 17, 1178616.

90. Kern, J.K., Geier, D.A., Sykes, L.K., and Geier, M.R. (2013). Evidence of neurodegeneration in autism spectrum disorder. Transl. Neurodegener. 2, 17.

91. Vivanti, G., Tao, S., Lyall, K., Robins, D.L., and Shea, L.L. (2021). The prevalence and incidence of early-onset dementia among adults with autism spectrum disorder. Autism Res. 14, 2189–2199.

92. Gallitano-Mendel, A., Izumi, Y., Tokuda, K., Zorumski, C.F., Howell, M.P., Muglia, L.J., Wozniak, D.F., and Milbrandt, J. (2007). The immediate early gene early growth response gene 3 mediates adaptation to stress and novelty. Neuroscience 148, 633–643.

93. Choi, E.Y., Franco, D., Stapf, C.A., Gordin, M., Chow, A., Cover, K.K., Chandra, R., and Lobo, M.K. (2023). Inducible CRISPR epigenome systems mimic cocaine induced bidirectional regulation of Nab2 and Egr3. J. Neurosci. 43, 2242–2259.

94. Chandra, R., Francis, T.C., Konkalmatt, P., Amgalan, A., Gancarz, A.M., Dietz, D.M., and Lobo, M.K. (2015). Opposing role for Egr3 in nucleus accumbens cell subtypes in cocaine action. J. Neurosci. 35, 7927–7937.

95. Marballi, K.K., Alganem, K., Brunwasser, S.J., Barkatullah, A., Meyers, K.T., Campbell, J.M., Ozols, A.B., Mccullumsmith, R.E., and Gallitano, A.L. (2022). Identification of activity-induced Egr3-dependent genes reveals genes associated with DNA damage response and schizophrenia. Transl. Psychiatry 12, 320.

96. Guo, A.-Y., Sun, J., Jia, P., and Zhao, Z. (2010). A novel microRNA and transcription factor mediated regulatory network in schizophrenia. BMC Syst. Biol. 4, 10.

97. Pfaffenseller, B., da Silva Magalhães, P.V., De Bastiani, M.A., Castro, M.A.A., Gallitano, A.L., Kapczinski, F., and Klamt, F. (2016). Differential expression of transcriptional regulatory units in the prefrontal cortex of patients with bipolar disorder: potential role of early growth response gene 3. Transl. Psychiatry 6, e805.

98. Canchi, S., Raao, B., Masliah, D., Rosenthal, S.B., Sasik, R., Fisch, K.M., De Jager, P.L., Bennett, D.A., and Rissman, R.A. (2019). Integrating gene and protein expression reveals perturbed functional networks in Alzheimer’s disease. Cell Rep. 28, 1103–1116.e4.

99. Juan, C.-X., Mao, Y., Han, X., Qian, H.-Y., and Chu, K.-K. (2024). EGR1 regulates SHANK3 transcription at different stages of brain development. Neuroscience 540, 27–37.

100. Tallafuss, A., Stednitz, S.J., Voeun, M., Levichev, A., Larsch, J., Eisen, J., and Washbourne, P. (2022). Egr1 is necessary for forebrain dopaminergic signaling during social behavior. eNeuro 9, ENEURO.0035–22.2022.

101. Velmeshev, D., Schirmer, L., Jung, D., Haeussler, M., Perez, Y., Mayer, S., Bhaduri, A., Goyal, N., Rowitch, D.H., and Kriegstein, A.R. (2019). Single-cell genomics identifies cell type-specific molecular changes in autism. Science 364, 685–689.

102. Saavedra-Bonilla, H., Varman, D.R., and Reyes-Haro, D. (2024). Spontaneous calcium transients recorded from striatal astrocytes in a preclinical model of autism. Neurochem. Res. 49, 3069–3077.

103. Séjourné, G., and Eroglu, C. (2024). Astrocyte-neuron crosstalk in neurodevelopmental disorders. Curr. Opin. Neurobiol. 89, 102925.

104. Allen, M., Huang, B.S., Notaras, M.J., Lodhi, A., Barrio-Alonso, E., Lituma, P.J., Wolujewicz, P., Witztum, J., Longo, F., Chen, M., et al. (2022). Astrocytes derived from ASD individuals alter behavior and destabilize neuronal activity through aberrant Ca2+ signaling. Mol. Psychiatry 27, 2470–2484.

105. Taliun, D., Harris, D.N., Kessler, M.D., Carlson, J., Szpiech, Z.A., Torres, R., Taliun, S.A.G., Corvelo, A., Gogarten, S.M., Kang, H.M., et al. (2021). Sequencing of 53,831 diverse genomes from the NHLBI TOPMed Program. Nature 590, 290–299.

106. Das, S., Forer, L., Schönherr, S., Sidore, C., Locke, A.E., Kwong, A., Vrieze, S.I., Chew, E.Y., Levy, S., McGue, M., et al. (2016). Next-generation genotype imputation service and methods. Nat. Genet. 48, 1284–1287.

107. 1000 Genomes Project Consortium, Auton, A., Brooks, L.D., Durbin, R.M., Garrison, E.P., Kang, H.M., Korbel, J.O., Marchini, J.L., McCarthy, S., McVean, G.A., et al. (2015). A global reference for human genetic variation. Nature 526, 68–74.

108. Kang, H.M., Subramaniam, M., Targ, S., Nguyen, M., Maliskova, L., McCarthy, E., Wan, E., Wong, S., Byrnes, L., Lanata, C.M., et al. (2018). Multiplexed droplet single-cell RNA-sequencing using natural genetic variation. Nat. Biotechnol. 36, 89–94.

109. Fleming, S.J., Chaffin, M.D., Arduini, A., Akkad, A.-D., Banks, E., Marioni, J.C., Philippakis, A.A., Ellinor, P.T., and Babadi, M. (2023). Unsupervised removal of systematic background noise from droplet-based single-cell experiments using CellBender. Nat. Methods 20, 1323–1335.

110. Hao, Y., Stuart, T., Kowalski, M.H., Choudhary, S., Hoffman, P., Hartman, A., Srivastava, A., Molla, G., Madad, S., Fernandez-Granda, C., et al. (2024). Dictionary learning for integrative, multimodal and scalable single-cell analysis. Nat. Biotechnol. 42, 293–304.

111. Korsunsky, I., Millard, N., Fan, J., Slowikowski, K., Zhang, F., Wei, K., Baglaenko, Y., Brenner, M., Loh, P.-R., and Raychaudhuri, S. (2019). Fast, sensitive and accurate integration of single-cell data with Harmony. Nat. Methods 16, 1289–1296.

112. Wu, T., Hu, E., Xu, S., Chen, M., Guo, P., Dai, Z., Feng, T., Zhou, L., Tang, W., Zhan, L., et al. (2021). clusterProfiler 4.0: A universal enrichment tool for interpreting omics data. Innovation (Camb.) 2, 100141.

113. Yu, G., Wang, L.-G., Han, Y., and He, Q.-Y. (2012). clusterProfiler: an R package for comparing biological themes among gene clusters. OMICS 16, 284–287.

114. Kanehisa, M., Furumichi, M., Tanabe, M., Sato, Y., and Morishima, K. (2017). KEGG: new perspectives on genomes, pathways, diseases and drugs. Nucleic Acids Res. 45, D353– D361.

115. Ashburner, M., Ball, C.A., Blake, J.A., Botstein, D., Butler, H., Cherry, J.M., Davis, A.P., Dolinski, K., Dwight, S.S., Eppig, J.T., et al. (2000). Gene ontology: tool for the unification of biology. The Gene Ontology Consortium. Nat. Genet. 25, 25–29.

116. Persad, S., Choo, Z.-N., Dien, C., Sohail, N., Masilionis, I., Chaligné, R., Nawy, T., Brown, C.C., Sharma, R., Pe’er, I., et al. (2023). SEACells infers transcriptional and epigenomic cellular states from single-cell genomics data. Nat. Biotechnol. 41, 1746–1757.

117. Stuart, T., Srivastava, A., Madad, S., Lareau, C.A., and Satija, R. (2021). Single-cell chromatin state analysis with Signac. Nat. Methods 18, 1333–1341.

118. Bulik-Sullivan, B.K., Loh, P.-R., Finucane, H.K., Ripke, S., Yang, J., Schizophrenia Working Group of the Psychiatric Genomics Consortium, Patterson, N., Daly, M.J., Price, A.L., and Neale, B.M. (2015). LD Score regression distinguishes confounding from polygenicity in genome-wide association studies. Nat. Genet. 47, 291–295.

